# Concordant transcriptional and morphological remodeling revealed by *in vivo* Perturb-CLEAR

**DOI:** 10.64898/2026.04.06.716787

**Authors:** Boli Wu, Sean K. Simmons, Seoyeon Kim, Jiwen Li, Masood A. Akram, Chang Sin Park, Xinhe Zheng, Iain Mendez, Sasha Patel, Alan Chau, Nadia Burciu, Pranay Dayal, Thokozile Nyasulu, Nhan Huynh, Grace S. Clarke, X. William Yang, Joshua Z. Levin, Xin Jin

## Abstract

The principle that form follows function has long guided thinking in biology and architecture alike. In the nervous system, however, form does more than reflect function: neuronal morphology actively constrains input patterns, synaptic integration, and circuit wiring. During postnatal neurodevelopment, dendritic architectures are assembled and remodeled through genetically encoded programs and activities, transforming molecular programs into circuit architecture. However, dendritic morphogenesis has been difficult to quantify at scale while systematically testing how genetic variants, including neurodevelopmental disorder (NDD) risk genes, alter these structures *in vivo*. We developed Perturb-CLEAR, which integrates pooled CRISPR screening and whole-mount imaging to quantify brain-wide cytoarchitecture, and paired it with Perturb-seq to link structural phenotypes to transcriptomic changes. Applying Perturb-CLEAR to the developing mouse cortex revealed morphogenesis trajectories accompanied by transcriptomic dynamics. Moreover, systematic perturbation of NDD risk genes uncovered gene-specific multimodal phenotypes. *Adnp* perturbation remodels basal dendrites in L4/5 IT (intratelencephalic) neurons but not other dendritic compartments or cell types, alongside consistent transcriptional shifts. Combined morphology and transcriptome analyses link NDD risk genes to concordant multimodal cellular phenotypes in the developing brain, highlighting diverse paths of perturbation effect propagation across modalities.

---

The mammalian brain comprises diverse cell types arranged into organized structures that support specific circuit activity and brain functions^1,2^. Neurons participate in these cytoarchitectures through their individual morphologies, with dendrite structure playing a key role in shaping neuronal activity. Long-range projections from distinct brain regions selectively target specific dendritic compartments, and these input-defined innervation patterns vary across postsynaptic cell types^3–5^. In the postsynaptic neuron, synaptic inputs across dendritic compartments are selectively tuned through location-dependent attenuation, nonlinear integration, and dendrite-soma coupling. Collectively, dendritic geometry directly impacts which inputs are received, how they are integrated, and transformed into spiking output^6,7^. Therefore, determining how dendritic compartments are built, and which genes that are responsible, is essential for linking molecular state to neuron function.

During postnatal development, cytoarchitecture is assembled through genetically encoded morphogenesis programs and refined by activity-dependent mechanisms^8–13^. Diverse gene families have been implicated in shaping neuronal form, including transcription factors, cytoskeletal components, and guidance molecules^14–17^. Despite this mechanistic foundation, we still lack a systematic *in vivo* understanding that links specific genes to dendrite compartment-specific remodeling, a key gap given that dendrites shape neuronal activity. Further, although multimodal studies link transcriptomic types to stereotyped morpho-electric phenotypes^18–22^, it remains unclear how neurons are shaped by transcriptional and morphological trajectories during development, and how neurodevelopmental disorder (NDD) risk genes shift those trajectories *in vivo*^23,24^. One clinically relevant context for these questions is the sensory cortex, given that atypical sensory reactivity is common across NDDs^25,26^. In the somatosensory cortex, thalamic sensory input is relayed through layer 4 and into layer 2/3 within cortical columns^27,28^. This well-defined laminar scaffold provides a tractable framework for interpreting compartment-specific dendritic remodeling and for linking genetic perturbations to circuit-relevant cellular phenotypes.

Recent technological advances have begun to address some aspects of this challenge. High-throughput CRISPR screens coupled with single-cell genomics (e.g., Perturb-seq) now enable massively parallel functional genomics *in vivo*^29–31^. In parallel, developments in spatial genomics and whole-mount imaging have made it possible to visualize brain-wide molecular or cellular organization^32,33^. However, three-dimensional (3D) neuronal morphology analysis platforms in intact tissue are not yet compatible with pooled, systematic genetic interrogation^21,34–40^. As a result, how genetic perturbations alter *in vivo* cytoarchitectural organization is typically examined one gene at a time^11,14,41^.

Here, we introduce Perturb-CLEAR (<u>C</u>ytoarchitecture and <u>L</u>ight-sheet <u>E</u>nabled <u>A</u>ssay of <u>R</u>eadouts), an *in vivo* phenotyping platform that combines sparse labeling, pooled genetic perturbations, and whole-mount light-sheet imaging for brain-wide cytoarchitecture mapping across scales. Integrating with Perturb-seq, we further connect structural to molecular phenotypes across development and genetic perturbation. Using this approach, we define dendritic compartment-specific morphogenesis trajectories and map how NDD risk gene perturbations reshape morphology and transcriptomic states *in vivo*.

## 3D cytoarchitecture mapping with single-cell resolution

To enable sparse single-neuron labeling, we performed *in utero* delivery of lentiviral vectors expressing a membrane-tethered tag^34,37^. We screened multiple variants and identified four tags, tdTomato, Myc, HA, and V5, that produced uniform cellular labeling (Extended Data Fig. 1A-B), with consistent morphometric measurements comparing tags (Methods and Extended Data Fig. 1C). We further optimized viral titers to achieve an optimal labeling density, yielding approximately 60 to 120 labeled neurons in the somatosensory cortex per brain while minimizing dendrite overlap (Methods and Extended Data Fig. 2A-B). Lentiviral vectors expressing membrane-tethered tdTomato (tdTomato-m) were injected into the lateral ventricles at embryonic day 13.5 (E13.5), which allows for transduction of progenitors and newly born neurons along the ventricular zone, achieving postnatal brain-wide labeling^29,31^. Brains were collected postnatally from P10 to P28 for morphological analysis (Methods, Fig. 1A, and Extended Data Fig. 3A). We obtained high-resolution, three-dimensional maps of labeled cells including both neurons and non-neuronal cells across major brain regions (Fig. 1B-C and Extended Data Fig. 3B-C). This approach allows for high-resolution phenotyping across scales: from subcellular and cellular structures, including dendritic spines and dendrites, to brain-wide structures, such as long-range axons (Extended Data Fig. 3D-E).

**Fig. 1.**
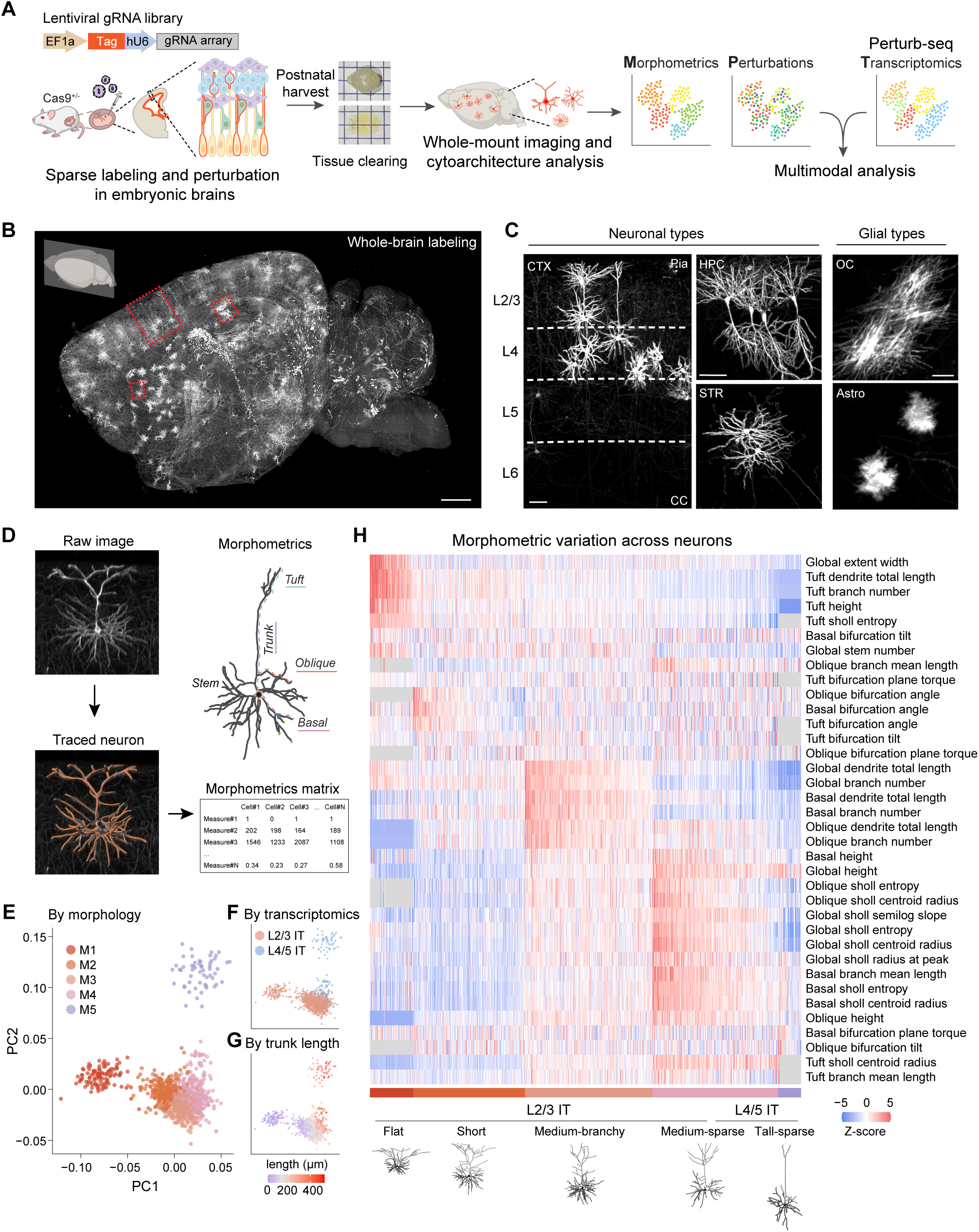
Perturb-CLEAR enables brain-wide cytoarchitecture mapping across cell types and reveals morphological diversity of upper layer cortical neurons. A. Schematics of Perturb-CLEAR, which combines *in vivo* sparse labeling, CRISPR editing and whole-mount imaging for cytoarchitectural mapping in developing brains. B. Representative light-sheet microscopy images (maximum projections of a hemisphere) of a cleared P21 brain sample injected with viral vectors expressing tdTomato-m and control gRNAs at E13.5. Scale bar: 1 mm. C. Representative light-sheet microscopy images (zoom-in from 1B) showing stereotypic cytoarchitectures of morphologically distinct cell types across brain regions. CTX, cortex; CC, corpus callosum; HPC, hippocampal complex; STR, striatum; OC, oligodendrocyte; Astro, astrocyte. Glial cell types were annotated visually by their stereotypic morphology. Scale bars: 100 μm. D. Schematic of cytoarchitectural analysis workflow. E. UMAP embeddings showing five morphological subtypes identified in upper layer somatosensory cortex. F. UMAP embeddings showing the inferred transcriptomic types by mapping morphology data to a public multimodal dataset^20^. G. UMAP embeddings showing the morphological diversity spectrum is correlated to trunk length of neurons, as a proxy for their cortical laminar locations. H. Heatmap and representative traces showing morphometric features of morphological subtypes. Morphometric measurements are normalized within each metric by z-score. NA values are in gray (Methods).

To systematically analyze the neuronal structural diversity in postnatal brains, we focused on pyramidal neurons from layer 2/3 and layer 4 of the somatosensory cortex, a population with stereotyped molecular signatures, anatomical layout, and circuit architecture (Methods and Extended Data Fig. 4A)^27,28,42^. To capture complete dendritic arbors, we detected membrane-tethered tags in cleared tissue, followed by segmentation, manual reconstruction, and rigorous quality control (Methods and Fig. 1D); throughout this study, we obtained 1,053 neuronal reconstructions (Extended Data Fig. 4B-C). For each neuron, we extracted a set of 170 morphometric features, including trunk length, dendritic extent, arborization complexity, branching angle, and Sholl profiles (Methods, Fig. 1D, Extended Data Fig. 5, Supplementary Table 1). These metrics captured major functionally distinct dendritic compartments that receive different inputs: tuft dendrites (long-range corticocortical inputs), oblique dendrites (local, feedforward signals), the apical trunk (tuft-soma coupling), and basal dendrites (local and ascending inputs)^6,7,43–46^. After removing the highly correlated metrics, we performed K-means clustering on 36 metrics and identified five major morphological subtypes (Methods and Fig. 1E). We then inferred the transcriptomic identities of these neurons by mapping morphometric profiles to a public multimodal dataset^20^, which assigned these neurons to L2/3 IT (intratelencephalic) or L4/5 IT neurons (Methods and Fig. 1F). Trunk length covaried with morphological subtype and inferred transcriptomic type (Fig. 1G), supporting the mapping accuracy and reinforcing the graded nature of cortical pyramidal neuron morphology^20,38,47^. For readability, we use the inferred transcriptomic labels L2/3 IT and L4/5 IT in later analyses, while noting that the reconstructed neurons were sampled anatomically from layer 2/3 and layer 4.

Across the five morphological subtypes (Fig. 1H and Extended Data Fig. 4D), superficial subtypes, M1 “Flat” and M2 “Short” types, showed smaller vertical extent and prominent tuft arborization. The intermediate classes, M3 “Medium-branchy” and M4 “Medium-sparse”, occupied overlapping laminar depths but differed continuously in oblique and basal dendritic complexity. The deepest class, M5, had the greatest overall height, consistent with its deeper position. All M-types were consistently observed across sample batches, confirming a robust cytoarchitectural spectrum across animals (Extended Data Fig. 4E). Together, these data define a morphometric spectrum of dendritic architecture in juvenile mouse somatosensory cortex, reinforcing the depth-associated graded variation described in other datasets^20,38,47^. This diversity may contribute to subtype-specific functional properties, including differences in local computation and long-range circuit connectivity^20,47–49^. We next examined the correspondence between morphological subtype and inferred transcriptomic identity. Across the five morphological subtypes, inferred L4/5 IT neurons were concentrated in M5 and part of M4, whereas inferred L2/3 IT neurons spanned M1-M4 (Fig. 1E-F). Our results support that transcriptomic identity broadly aligns with morphology, but the correspondence is graded rather than one-to-one at finer resolution^20,47^.

## Concordant morphogenesis across morphological and transcriptomic modalities

We next examined how this architecture is refined during postnatal development. Prior work has shown that postnatal dendritic and synaptic maturation unfolds over discrete developmental windows, and that these dynamics can vary across cortical layers, areas, and dendritic compartments^8,50–55^. However, it remains unclear how postnatal structural changes relate to developmental gene-expression programs. We therefore applied Perturb-CLEAR to quantify developmental changes in dendrites and dendritic spines (as a structural proxy of synapses^56^) of layer 2/3 and layer 4 cortical pyramidal neurons in the somatosensory cortex and related these structural dynamics to transcriptional programs from a public atlas across P10, P14, and P28 (Fig. 2A).

**Fig. 2.**
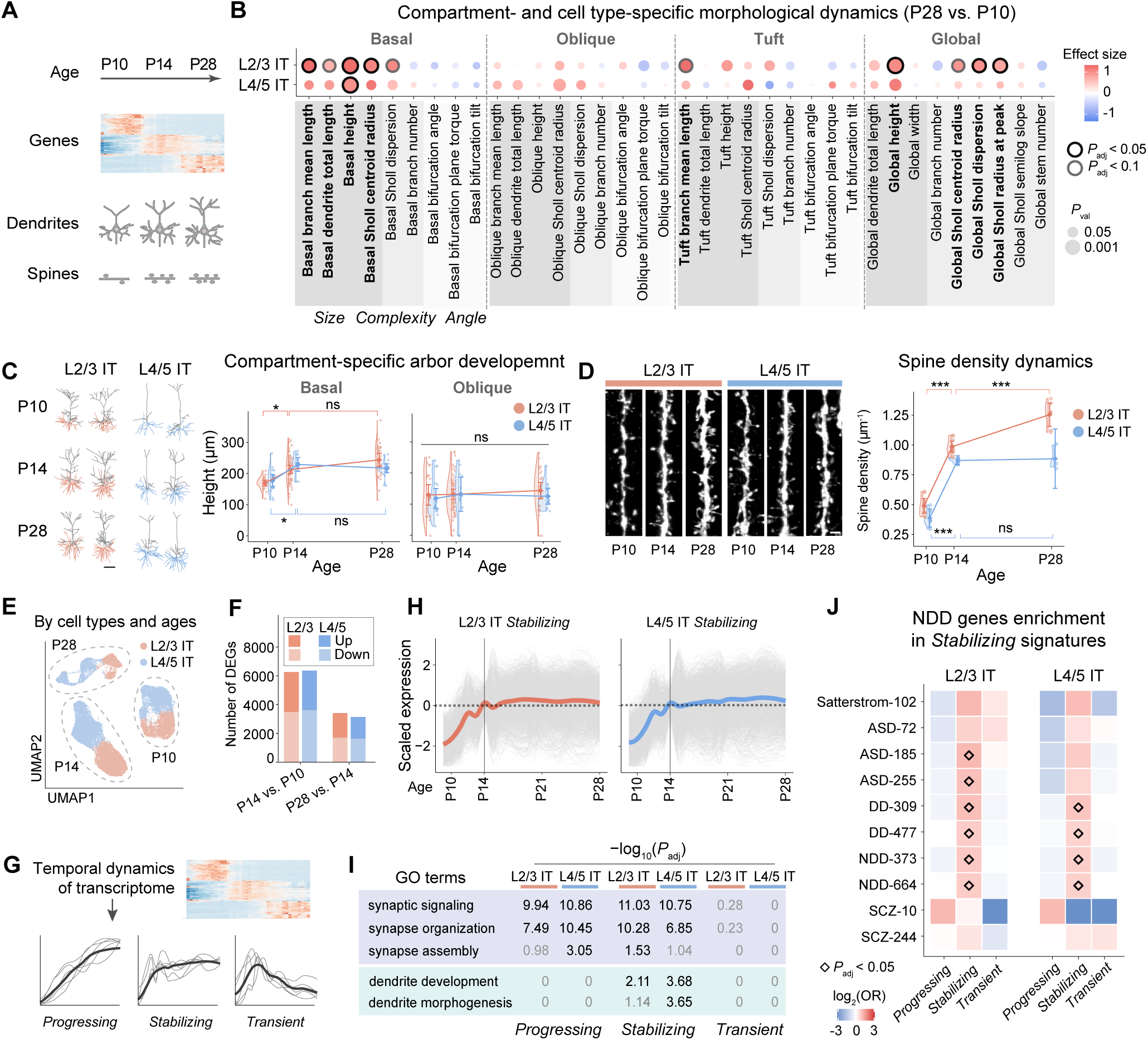
Dendritic compartment- and cell type-specific morphogenesis dynamics across development with concordant transcriptomic signatures. A. Schematics of developmental neuronal changes across modalities: gene expression, dendrites, and spines (as a structural proxy of synapses). B. Dot plot showing shared and distinct morphometrics with significant differences comparing P28 to P10, across wild-type L2/3 IT and L4/5 IT neurons. Different gray shadings indicate categories of morphometric features (size, complexity, and angle). P10: n = 21 L2/3 IT neurons from 4 brains, n = 17 L4/5 IT neurons from 4 brains. P28: n = 28 L2/3 IT neurons from 4 brains, n = 14 L4/5 IT neurons from 3 brains. C. Representative neuron traces and plot showing dendritic compartment-specific arbor development dynamics across three ages (P10, P14, and P28). Basal compartments were marked in orange (L2/3 IT) or blue (L4/5 IT) in the raw traces. Metrics were plotted over numeric postnatal ages, with conditions offset slightly for visualization purposes. Raincloud plots show half violins and half scattered points (individual neurons). Diamonds mark group means, and error bars represent t-based 95% confidence interval. *: *P*_adj_ < 0.05; ns: *P*_adj_ > 0.05. Scale bar: 100 μm. D. Representative images and quantification of developmental dynamics of spine density across ages (P10, P14, and P28) and cell types (L2/3 IT and L4/5 IT). Data were plotted over numeric postnatal ages, with conditions offset slightly for visualization purposes. Raincloud plots show half violins and half scattered points (individual neurons). Diamonds mark group means, and error bars represent t-based 95% confidence interval. P10: n = 9 L2/3 IT neurons from 4 brains, n = 5 L4/5 IT neurons from 2 brains; P14: n = 9 L2/3 IT neurons from 2 brains, n = 4 L4/5 IT neurons from 2 brains; P28: n = 7 L2/3 IT neurons from 2 brains, n = 3 L4/5 IT neurons from 1 brain. ***: *P*_adj_ < 0.001; ns: *P*_adj_ > 0.05. E. UMAP embedding showing cell type clustering across ages^59^, n = 20,614 L2/3 IT neurons, n = 26,538 L4/5 IT neurons. F. Bar plot showing differentially expressed gene (DEG) number across ages in both cell types (*P*_adj_ < 0.01). P14 vs. P10: L2/3 IT, 2790 up, 3472 down; L4/5 IT, 2762 up, 3602 down. P28 vs. P14: L2/3 IT, 1718 up, 1694 down; L4/5 IT, 1526 up, 1610 down. G. Schematics showing gene expression analysis to identify three gene signatures (*Progressing*, *Stabilizing*, and *Transient*) based on temporal dynamics of gene expression. H. Developmental expression trajectories of genes within the *Stabilizing* signatures. Scaled expression level relative to P14 time point is shown. L2/3 IT: n = 1963 genes; L4/5 IT: n = 2185 genes. I. Gene ontology analysis shows significant enrichment of synapse-related terms (signaling and structure) in the *Progressing* and the *Stabilizing* signatures for both cell types, while dendrite-related morphogenesis terms are only enriched in the *Stabilizing* signatures. Black: *P*_adj_ significant. Gray: *P*_adj_ non-significant. J. Fisher exact tests reveal significant enrichment of ASD, NDD, and DD de novo risk genes specifically in the *Stabilizing* signatures of both cell types, but not other signatures. ASD: autism spectrum disorders. DD: developmental delay. NDD: neurodevelopmental disorders. SCZ: schizophrenia.

We focused on the 144 wild-type neuron reconstructions from the dataset across cell types (inferred L2/3 IT and L4/5 IT neurons) and ages (P10, P14, and P28) (Extended Data Fig. 6A-B and Supplementary Table 2) and revealed cell type- and dendritic compartment-specific changes (Fig. 2B and Extended Data Fig. 7A-C). The largest dendritic changes were observed in the basal arbor (Fig. 2B). Basal dendrites expanded in both L2/3 IT and L4/5 IT neurons (e.g., L2/3 basal height, P28 vs. P10: effect size = 1.45, *P*_adj_ = 3.63×10^−6^; Fig. 2B), whereas tuft and oblique dendrites were comparatively stable over the same period (Fig. 2B and Extended Data Fig. 7A). The intermediate time point (P14) further localized most of this change to an early phase: the largest changes occurred from P10 to P14, followed by a stabilization phase from P14 to P28 (Fig. 2C and Extended Data Fig. 6C). We next assessed basal dendritic spine density as a proxy for excitatory synapse abundance in Perturb-CLEAR-labeled neurons across the same three ages, which showed a similar stabilizing trajectory in both L2/3 IT and L4/5 IT neurons (Methods, Fig. 2D and Supplementary Table 3). Thus, dendritic growth and spine accumulation followed similar temporal trends in early postnatal somatosensory IT neurons.

Together, our results from these wild-type neurons reinforce a model in which postnatal maturation is temporally structured, dendritic compartment-specific, and cell type-specific rather than uniform^8,50,53^. Within this framework, the largest dendritic changes in these somatosensory IT neurons occur in the basal compartment and that basal spine density follows a similar temporal trend. The more prolonged structural maturation observed in L2/3 IT neurons further echoes prior evidence that, in somatosensory cortex, layer 2/3 circuits remain plastic over a longer postnatal interval, whereas layer 4 circuits finish maturation earlier^57,58^. The similar temporal trends of dendrite and spine dynamics are consistent with coordinated structural maturation during cortical circuit formation and refinement.

To relate postnatal morphogenesis to transcriptional dynamics, we next leveraged a public developmental transcriptomic atlas with dense temporal sampling across early postnatal maturation^59,60^ (Methods and Fig. 2E). In both L2/3 IT and L4/5 IT neurons, we identified differentially expressed genes (DEGs) between ages (Fig. 2F and Supplementary Table 4). To identify gene signatures correlated with morphogenesis, we grouped genes that were upregulated at P14 relative to P10 by their subsequent expression at P28 into three signatures: *Progressing*, *Stabilizing*, and *Transient* (Fig. 2G-H, Extended Data Fig. 8A-B and Supplementary Table 5). Notably, gene set enrichment analysis revealed that synaptogenesis-related terms are preferentially enriched among the *Progressing* and *Stabilizing* signatures across cell types, whereas dendritogenesis-related terms are selectively enriched in the *Stabilizing* signature (Fig. 2I and Supplementary Table 6). This is consistent with the early growth and later stabilization dynamics observed in morphology data (Fig. 2B-C). In addition, autism spectrum disorder (ASD) and broader NDD risk genes are selectively enriched in the *Stabilizing* signatures in both cell types (Fig. 2J and Supplementary Table 6)^61–64^. These results support aligned developmental trajectories and time courses in morphology and transcriptome, pointing to an early postnatal window as a candidate epoch of NDD vulnerability. We therefore asked how NDD risk genes perturb both modalities during this period.

## NDD risk genes drive morphological remodeling in distinct compartments and cell types

We applied Perturb-CLEAR to interrogate neuronal morphological changes following perturbation of a panel of ten NDD risk genes (developmental expression levels in Extended Data Fig. 9A). We focused on genes encoding synaptic proteins (e.g., *Grin2b*, *Syngap1*) with established roles in dendrite or synapse maturation^65–67^, alongside transcription factors or chromatin remodelers (e.g., *Adnp*, *Foxp1*) that may influence morphology through broader regulatory programs^68,69^. We generated a pooled gRNA lentiviral library targeting early coding exons of the genes (Methods and Supplementary Table 7). After validating gRNA efficiency *in vitro* and quality control of the gRNA library (Methods, Extended Data Fig. 9B-C), we performed embryonic injection of the library at E15.5, enabling mosaic perturbations in layer 2/3 and layer 4. Each perturbation or control was barcoded by a unique fluorescent tag or tag combination that was optically distinguishable (Methods, Fig. 3A, and Extended Data Fig. 9D). Viral titers were assessed to achieve low multiplex labeling to maximize single perturbations (Extended Data Fig. 2C-D).

**Fig. 3.**
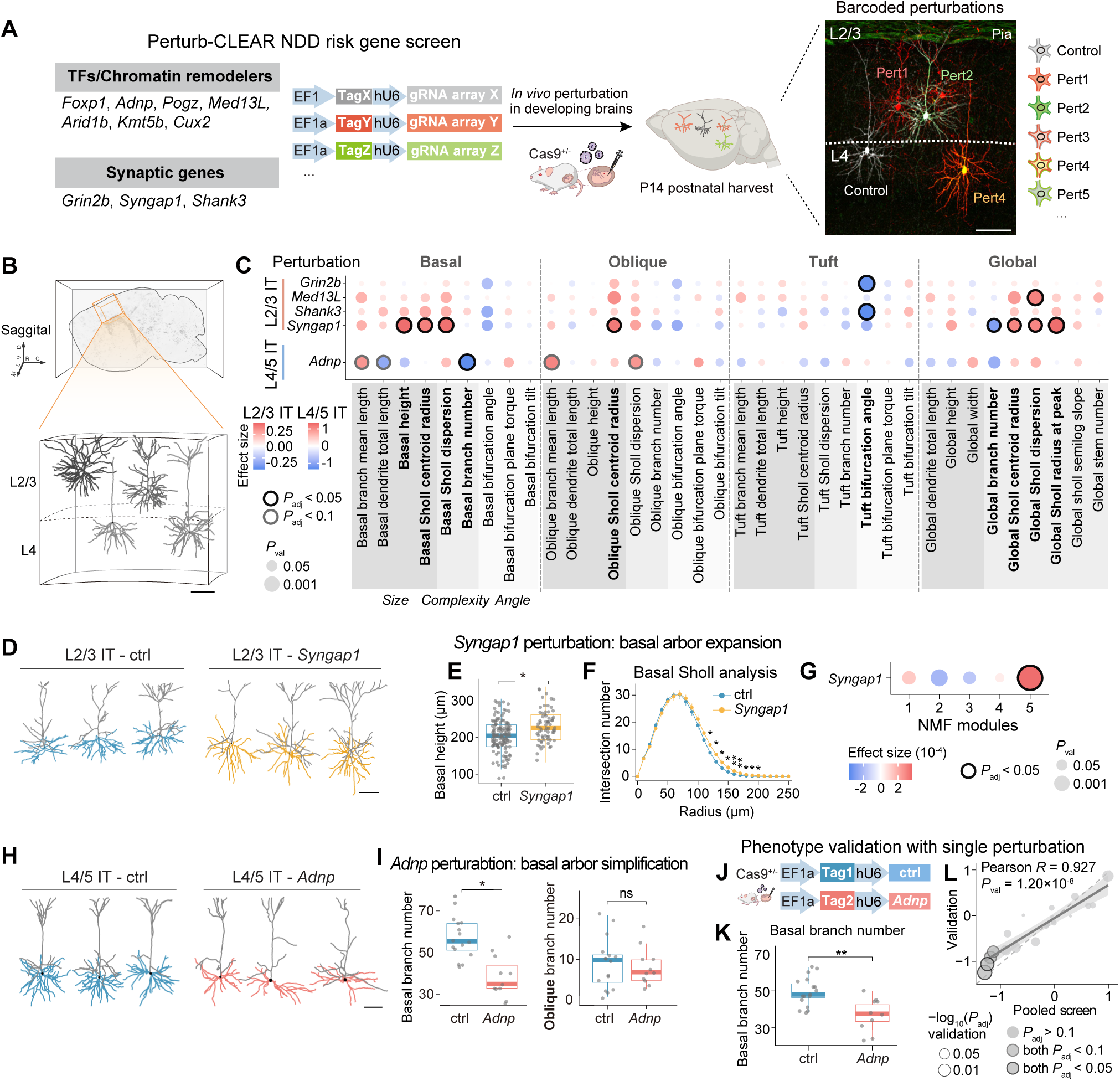
NDD risk gene perturbation leads to cell type- and dendritic compartment-specific morphology changes. A. Schematics of NDD risk gene screen. Representative immunostaining image shows multiplex labeling in layer 2/3 and layer 4 somatosensory cortex, with each tag variant corresponding to one perturbation or control. TFs, transcription factors. Scale bar: 100 μm. B. Schematics of 3D morphology reconstruction of neurons in upper layer somatosensory cortex in whole-mount tissue. Scale bar: 100 μm. C. Dot plot showing specific morphometric changes elicited by NDD risk gene perturbations across cell types and perturbations. Different gray shadings indicate categories of morphometric features (size, complexity, and angle). n ≥ 60 neurons from N ≥ 9 brains per perturbation for L2/3 IT neurons; n ≥ 10 neurons from N ≥ 5 brains per perturbation for L4/5 IT neurons. Sample details were included in Supplementary Table 8. D. Representative traces showing *Syngap1* perturbation leads to basal dendrite arborization expansion in L2/3 IT neurons. Basal compartments were marked in blue (control) or yellow (*Syngap1*-perturbed) in the raw traces. Scale bar: 100 μm. E. Boxplots showing significant increase of basal height in *Syngap1* perturbed L2/3 IT neurons. The interquartile range (IQR) with median (center line) is shown, with whiskers extending to 1.5× IQR; overlaid jittered points indicate individual neurons. *Syngap1-*perturbed: n = 68 neurons from 9 brains, control: n = 135 neurons from 19 brains. *: *P*_adj_ < 0.05. F. Sholl analysis of basal dendrites from L2/3 IT neurons. Data are shown as mean ± SEM. Group differences at each radius were assessed by two-sided Wilcoxon rank-sum tests, with neurons treated as replicates, followed by Benjamini-Hochberg correction across radii. *: *P_a_*_dj_ < 0.05; **: *P*_adj_ < 0.01. G. Non-negative matrix factorization module analysis showing specific enrichment of module 5 in *Syngap1*-perturbed neurons. H. Representative traces showing *Adnp* perturbation leads to simplification of the basal dendrite arborization in L4/5 IT neurons. Basal compartments were marked in blue (control) or red (*Adnp*-perturbed) in the raw traces. Scale bar: 100 μm. I. Boxplots showing significant decrease of basal branch number of *Adnp* perturbed L4/5 IT neurons, but not oblique branch number. The interquartile range (IQR) with median (center line) is shown, with whiskers extending to 1.5× IQR; overlaid jittered points indicate individual neurons*. Adnp-*perturbed: n = 10 neurons from 5 brains, control: n = 27 neurons from 10 brains. *: *P*_adj_ < 0.05; ns: *P*_adj_ > 0.05. J. Schematics of single perturbation experiment, where *Adnp* and control gRNAs are introduced into the developing brain for postnatal harvest at P14. K. Boxplots of *Adnp* perturbed L4/5 IT neurons showing decreased basal branch number in single perturbation validation. The interquartile range (IQR) with median (center line) is shown, with whiskers extending to 1.5× IQR; overlaid jittered points indicate individual neurons. *Adnp*-perturbed: n = 10 from 3 brains, control: n = 17 from 6 brains. *: *P*_adj_ < 0.05. L. Phenotypic effect size of basal compartment-related metrics is correlated between pooled screen and single validation dataset. A fitted linear regression line (solid) is shown with 95% confidence band. Dashed line indicates y = x.

In total, we collected 909 high-quality reconstructions from neurons sampled in layer 2/3 and layer 4 of somatosensory cortex in whole-mount cleared tissue, with an average of 82 cells assayed per perturbation (Fig. 3B, Extended Data Fig. 9E and Supplementary Table 8). After mapping neurons to transcriptomic cell types (L2/3 IT and L4/5 IT), we observed no significant perturbation-driven shifts in inferred transcriptomic cell type proportions (Extended Data Fig. 9F)^20^. Further, morphometric analysis identified perturbation-specific structural remodeling from five genes across basal, oblique, and tuft compartments (*Grin2b, Med13L, Shank3, Syngap1,* and *Adnp*) (Methods, Fig. 3C, Extended Data Fig. 10 and Supplementary Table 9).

*Syngap1* perturbation led to one of the strongest phenotypes, showing selective expansion of the basal dendritic arborization in L2/3 IT but not in L4/5 IT neurons (Fig. 3C-G and Extended Data Fig. 11A-C). We observed a significant increase in basal height (effect size = 0.49, *P*_adj_ = 3.45×10^−2^) and a rightward shift in Sholl intersection profiles (Fig. 3E-F), as well as a set of other basal and global-related metrics (including global Sholl radius at peak, basal Sholl centroid radius, and basal Sholl dispersion, Extended Data Fig. 11A). Notably, not all arborization size-related metrics change in the concordant direction: while most metrics indicate an arborization expansion in *Syngap1*-perturbed neurons, global branch number exhibited a decrease trend (Extended Data Fig. 11A), highlighting the necessity of comprehensive morphometric profiling. To test whether these metrics reflect a coordinated remodeling program, we performed non-negative matrix factorization (NMF) to extract sets of covarying metrics (“NMF modules”) and quantified each neuron’s module score (Methods, Fig. 3G and Extended Data Fig. 11D-E). Among five modules identified, NMF module 5 was preferentially elevated in *Syngap1*-perturbed L2/3 IT neurons, impacting several metrics collectively (Fig. 3G and Extended Data Fig. 11E). These data indicate that *Syngap1* perturbation has coordinated effects on dendritic architecture rather than affecting individual metric independently, supporting *Syngap1*’s role as a regulator of dendrite and synapse maturation during development^66,67,70,71^.

*Adnp* perturbation in L4/5 IT neurons also resulted in significant cytoarchitectural changes, leading to simplification of basal dendrites (Fig. 3H). Interestingly, this phenotype is highly dendritic compartment-specific, with significant decreases in basal branches (effect size = −1.29, *P*_adj_ = 1.77×10^−2^) while oblique branches are largely unaffected (Fig. 3I). To validate this result from the pooled screen, we introduced *Adnp*-targeting and control gRNAs into developing brains (Fig. 3J and Supplementary Table 8). Consistently, we observed a reduction in basal branch number within the same neuronal population (Fig. 3K); the effect sizes for other basal-compartment metrics were highly correlated with those of the pooled screen (Pearson *R* = 0.927, *P*_val_ = 1.20×10^−8^) (Fig. 3L, Extended Data Fig. 11F-G and Supplementary Table 10). Together, these data demonstrate that NDD risk genes reshape development through selective vulnerability of specific neuronal cell types and dendritic compartments that can be resolved robustly with Perturb-CLEAR.

## NDD risk genes orchestrate cell type-specific transcriptional shifts

To connect these structural phenotypes to underlying molecular programs, we profiled a matched perturbation panel at the same developmental stage using *in vivo* Perturb-seq^31^. Adeno-associated viruses (AAVs) expressing a nucleus-anchoring GFP and gRNAs were pooled and injected at day E15.5 (Fig. 4A and Extended Data Fig. 12A-B). At P14, we lysed the neocortex and enriched GFP^+^ nuclei for 5′ single-nucleus RNA-seq (snRNA-seq) (Extended Data Fig. 12C-D). After quality control, we retrieved 50,846 high-quality nuclei assigned with high-confidence perturbation identities from 11 animals across 4 batches for phenotypic analysis (Methods, Fig. 4A, Extended Data Fig. 12E-K, and Supplementary Table 11); perturbations were represented by a median of 2,654 nuclei each. We retained 10 annotated cell types for downstream analysis: eight clusters of excitatory glutamatergic neurons (L2/3 IT, L4/5 IT, L5 IT, L6 IT, L5 PT (pyramidal tract), L5 NP (near-projecting), L6 CT (corticothalamic), and L6b) and two clusters of GABAergic inhibitory neurons (Sst and Pvalb) (Methods and Fig. 4B)^42^. We did not detect major cell type proportion changes resulting from the perturbations, consistent with the Perturb-CLEAR results (Extended Data Fig. 13).

**Fig. 4.**
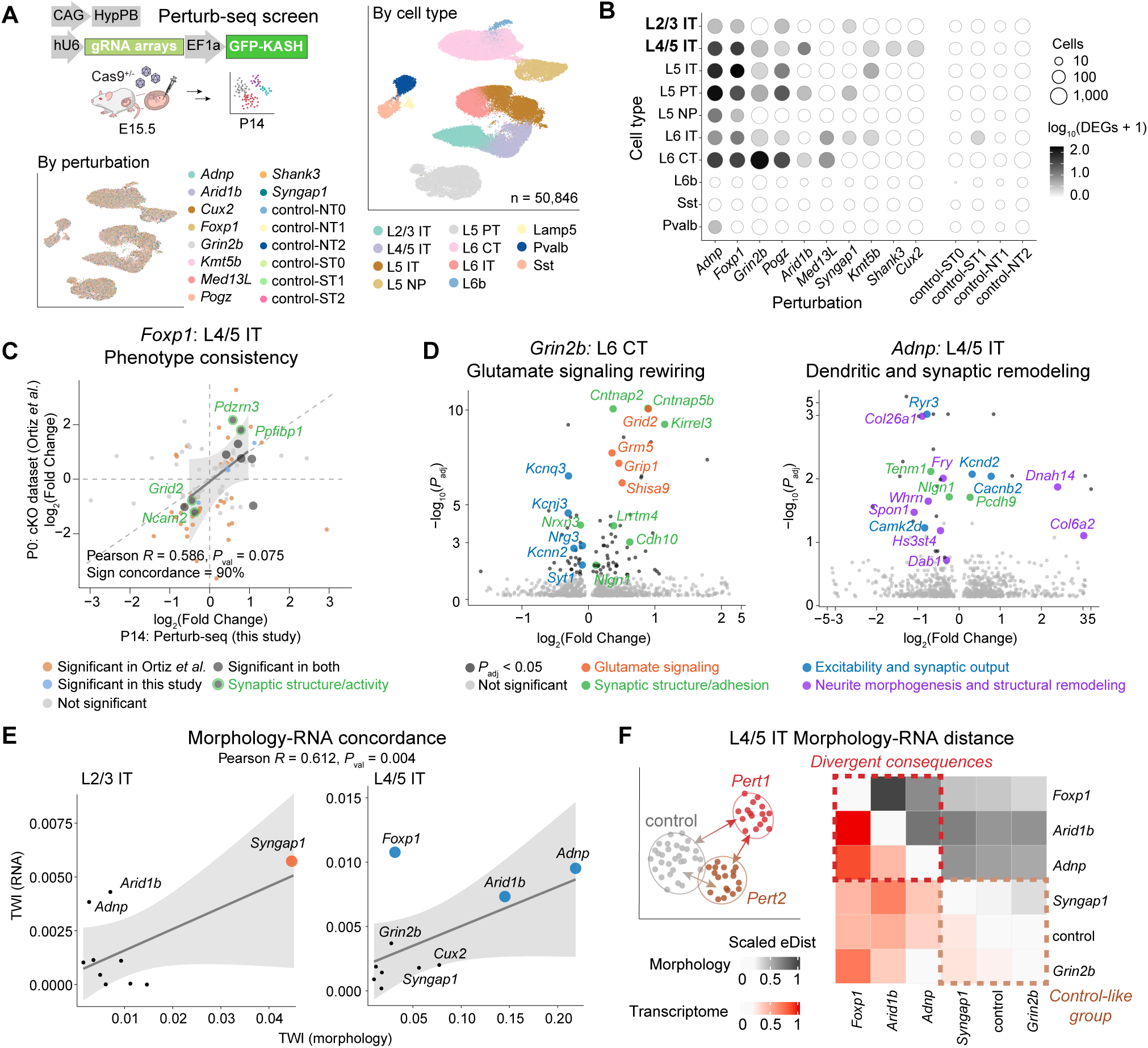
Joint analysis of NDD risk gene Perturb-seq and Perturb-CLEAR screens uncovers concordant changes of genes and morphology. A. Upper left: Schematics of *in vivo* Perturb-seq of NDD risk genes, which introduces pooled perturbations in the developing brain by *in utero* AAV-SCH9 injection. The phenotype is examined postnatally at P14 through single-nucleus RNA-seq using fluorescence-activated nuclei sorting (FANS)-purified perturbed nuclei. Right: UMAP embedding showing 50,846 high-quality, single perturbation cells clustered by cell types. Lower left: UMAP embedding showing cell clusters colored by assigned perturbation identities. B. Dot plot showing number of DEGs identified by Perturb-seq analysis. C. Correlation of the DEG log_2_(Fold Changes) derived from *Foxp1* perturbed L4/5 IT neurons from Perturb-seq data in this study and those from *Foxp1* cKO L4/5 IT neurons from a public dataset^72^. All shared-nominal DEGs are shown. Only shared-FDR significant DEGs were used for correlation calculation and sign concordance. A fitted linear regression line (solid) is shown with 95% confidence band. Dashed lines indicate x = 0, y = 0, and y= x. D. Volcano plot showing perturbation- and cell type-specific DEGs. Left: *Grin2b* perturbed L6 CT neurons showing DEGs enriched in synaptic structure/adhesion, glutamate signaling, as well as neuron excitability; Right: *Adnp* perturbed L4/5 IT neurons showing DEGs enriched in synaptic structure, neuron excitability, and neurite morphogenesis and structural remodeling programs. E. Correlation of transcriptome wide impact (TWI) that measures perturbation-induced phenotypic effect size for both morphology and RNA. For each data point, percentages relative to the maximum were calculated separately within each measurement and then averaged; points exceeding 20% were labeled, and those exceeding 20% were displayed at a larger size. A fitted linear regression line is shown with 95% confidence band for each cell type. F. Cross-modality distance of morphology and RNA in L4/5 IT neurons. Upper-left: schematics of energy distance (eDist) analysis measuring perturbation-control distance and pairwise distance between perturbations. Right: Heatmap of eDist showing diverse phenotypic effects across perturbations in the morphological modality (black) or transcriptomic modality (red). Perturbations with n < 5 neurons sampled were removed.

We compared the cell type-specific transcriptomes in perturbed cells to control cells (control-ST2) (Methods). Four perturbations produced over 20 DEGs in subsets of cell types (Fig. 4B and Supplementary Table 12): *Adnp* and *Foxp1* induced DEGs across cell types (notably L4/5 IT, L5 IT, L5 PT, and L6 CT), whereas *Pogz* and *Grin2b* produced strong gene expression shifts specifically in L6 CT. Control comparisons (e.g., control-NT2) yielded 0-1 DEGs, indicating low false-positive discoveries.

We first investigated the phenotype of *Foxp1*, which produced a widespread impact on synaptic structure and adhesion modules across cell types (*P_adj_* < 0.05, L4/5 IT: 34, L5 IT: 114, L5 PT: 22, and L6 CT: 29 DEGs, Extended Data Fig. 14A). We compared our phenotypes with a published *Foxp1* cKO scRNA-seq dataset^72^ and observed correlated log_2_(Fold Changes) of shared DEGs (Supplementary Table 13) in L4/5 IT (90% sign concordance; Pearson *R* = 0.586, *P*_val_ = 7.53×10^−2^; Fig. 4C) and L5 PT (Extended Data Fig. 14B). Specifically, we found concordant changes in synaptic structure and activity remodeling genes, including *Pdzrn3, Grid2*, etc., supporting that *Foxp1* impacts synaptic function^68,73^.

Among the most pronounced and selective phenotypes, *Grin2b* perturbation elicited a large transcriptional response in L6 CT (93 DEGs, *P*_adj_ < 0.05; Fig. 4D left), with minimal DEGs in all other major cell types (Extended Data Fig. 14C). We first observed a predominantly upregulated gene program related to synaptic structure (*Kirrel3, Cntnap2, Nlgn1*) (Fig. 4D left), accompanied by upregulation of genes that may indicate compensatory glutamatergic signaling, including AMPA-associated synaptic components (*Lrrtm4*, *Grip1*)^65,74–77^ and other glutamate receptors/regulators (*Grm5, Grid2*, *Shisa9*)^78–80^. In addition, we observed a coherent reduction of potassium channel (*Kcnq3, Kcnj3, Kcnn2*) and vesicle release-related genes (*Nrg3, Syt1*)^81,82^, suggesting that *Grin2b* loss may impact intrinsic excitability and synaptic output^83^. Collectively, these transcriptional changes align with NDD frameworks implicating disrupted synaptic transmission and E/I homeostasis^84^, while also pointing to an L6 CT-specific response that is particularly relevant for corticothalamic control of sensory throughput at this age. Given evidence linking atypical thalamocortical interactions to sensory sensitivities and symptom severity in NDD^25,26,85,86^, this result provides a plausible molecular entry point for disease-associated alterations in sensory processing, while the circuit consequences remain to be tested directly.

We next examined *Adnp*, an NDD risk gene with reported roles spanning transcriptional regulation and cytoskeleton modulation^69,87–90^. Within L4/5 IT neurons, *Adnp* perturbation DEGs were enriched for programs related to synaptic structure, neuronal excitability, and structural remodeling, including ECM organization (Fig. 4D right, *Col6a2*, *Col26a1*, etc.) and cytoskeleton regulation (*Fry*, *Dnah14*, etc.). These transcriptional changes are consistent with the basal dendrite simplification observed in L4/5 IT neurons in the Perturb-CLEAR morphology data (Fig. 3H-L). By contrast, L2/3 IT lacked overt morphological changes, underscoring the cell type-specific translation of transcriptional perturbation into structural outcomes (Extended Data Fig. 10A and Extended Data Fig. 14D). Together, these findings demonstrate that NDD risk genes engage cell type-specific molecular programs coupled to dendritic compartment-specific cytoarchitectural remodeling.

## Morphology-RNA concordance and discordance

These paired datasets allowed us to compare perturbation effects in two complementary ways: their overall magnitude of deviation from the controls, and whether different perturbations shift cells toward similar phenotypic states. We first assessed effect magnitudes in transcriptomic and morphometric space using transcriptome wide impact (TWI) relative to controls as a summary measure (Methods, Fig. 4E and Supplementary Table 14). Across cell types, perturbations with larger transcriptomic shifts generally also show larger morphological shifts, indicating broad concordance between molecular and structural phenotypes (Pearson *R* = 0.612, *P*_val_ = 4.14×10^−3^). Several NDD genes, including *Syngap1* in L2/3 IT neurons and *Adnp* and *Arid1b* in L4/5 IT neurons, showed concordant effects across modalities. By contrast, *Foxp1* in L4/5 IT neurons displayed strong transcriptomic changes but minimal morphological changes at this age, suggesting dissociable molecular and structural responses.

We next asked whether perturbations with similar effect magnitudes also produced similar phenotypes. To address this, we quantified pairwise energy distance (eDist) among perturbations, as well as between each perturbation and control, where lower values indicate greater similarity (Methods, Fig. 4F, Extended Data Fig. 15A, and Supplementary Table 14). This analysis first recapitulated the control-distance patterns observed with TWI (Fig. 4E): e.g., *Syngap1* and *Grin2b* showed minimal transcriptional deviation from control in L4/5 IT neurons (*Syngap1*-control_RNA_ = 0.245; *Grin2b*-control_RNA_ = 0.200) (Fig. 4F). Further, eDist showed that perturbations did not drive a shared phenotype, but instead displaced cells heterogeneously from controls and from one another across modalities (e.g., *Foxp1*, *Adnp*, and *Arid1b* in L4/5 IT neurons). *Adnp* and *Arid1b* provide an example: despite similar phenotypic strength across modalities (Fig. 4E), they remained separated by a large eDist (*Adnp*-*Arid1b*_RNA_ = 0.443; *Adnp*-*Arid1b*_Morphology_ = 0.767), showed limited overlap at the DEG level (Fig. 4D and Extended Data Fig. 15B), nor do they produce the same morphological outcomes (Extended Data Fig. 10). Together, these results show that despite broad concordance in overall perturbation magnitudes, similar perturbation magnitudes do not imply shared phenotypic trajectories. Instead, NDD gene perturbations can elicit diverse cellular responses across modalities, underscoring the value of joint profiling.

## Discussion

In this study, we paired Perturb-CLEAR with matched Perturb-seq to relate neuronal morphology to transcriptomic state *in vivo*. This framework resolved compartment-specific morphogenesis trajectories in wild-type neurons across development (Fig. 2B) and, with genetic perturbations, revealed gene- and cell type-specific phenotypes across dendritic architecture (Fig. 3C) and transcriptional changes (Fig. 4B). Notably, coupling across morphology and transcriptome, together with context-dependent decoupling (Fig. 4E-F), emerged as a central feature of the perturbation response. Prior multimodal atlases established that transcriptomic identity constrains broad morpho-electric phenotype, rather than one-to-one at finer scales^18–21^. Our results extend this framework into a perturbation setting and delineate how morphology-transcriptome correspondence can be differentially engaged and remodeled by disease-relevant genetics.

When transcriptomic and morphological phenotypes were concordant, we interpret this as evidence that a perturbation engaged molecular programs that propagated to implemented structural remodeling *in vivo*. Apparent discordance could be interpreted in different ways: it may reflect mechanisms including buffering or compensation, or a temporal offset between early molecular responses and later structural changes. It may also reflect regulatory layers that are incompletely captured by snRNA-seq^91^, including intrinsic or extrinsic cues, circuit activities, or synaptic and cytoskeleton regulators^14–17^. *Adnp*-perturbed L4/5 IT neurons and *Syngap1*-perturbed L2/3 IT neurons illustrate this principle. Both showed propagated phenotypes across modalities, yet they culminated in distinct cell type-specific and compartment-specific dendritic alterations (Fig. 3C-I). As neuronal morphology actively constrains input patterns and synaptic integration: these dendritic phenotypes provide plausible hypotheses of how NDD risk gene perturbations may influence circuit assembly in somatosensory processing, though their functional consequences remained to be tested directly. Collectively, we highlight that NDD perturbations may influence neuron function through distinct molecular and cellular implementations rather than through a single shared route. Perturbational phenotypes are thereby hard to predict, either from effect magnitude alone, or from one modality to another. Future screens that integrate same-cell multimodal measurements across development, ideally with circuit-level readouts, larger screening scale to include more gene panels, will be important for tracing how perturbation effects propagate, diverge, and converge during brain development. Taken together, our results argue that mechanistic interpretation of disease genetics requires tracking perturbation effects across modalities, rather than inferring outcome from a single readout alone.

## Methods

### Animals

All animal experiments were performed according to protocols approved by the Institutional Animal Care and Use Committees (IACUC) of The Scripps Research Institute. All mice were kept in standard conditions (a 12-hour light/dark cycle with ad libitum access to food and water). Constitutively expressing Cas9 (Jax#026179) and wild-type mice (JAX #000664) were obtained from Jackson Labs.

### Mammalian cell culture

Mammalian cell culture experiments were performed in the HT22-Cas9 mouse hippocampal neuronal cell line (Ubigene, YC-A004-Cas9-H) or HEK293T cell line (ATCC, #CRL-3216) grown in DMEM (Thermo Fisher Scientific, #11965092) with 25 mM glucose, 1 mM sodium pyruvate and 4 mM L-Glutamine (Thermo Fisher Scientific, #11995073), additionally supplemented with 1% penicillin–streptomycin (Thermo Fisher Scientific, #15140122), and 5-10% fetal bovine serum (Thermo Fisher Scientific, #16000069). HT-22 cells were maintained at confluency below 80%. Transfections were performed with PEI (Polysciences, #24765-1) or Lipofectamine™ 3000 Transfection Reagent (Thermo Fisher Scientific, L3000001).

### gRNA design and editing efficiency tests

gRNA design was performed as previously described^24^. In brief, the gRNAs were designed using an online tool (benchling.com), gRNAs were designed for each gene targeting the early coding exons of the transcript. Control gRNAs (including six safe-targeting and six non-targeting controls) were adopted from a published gRNA library^92^. The full sequences of the gRNAs used in this work are listed in Supplementary Table 7. For in vitro knockout efficiency validation, HT22-Cas9 (Ubigene, YC-A004-Cas9-H) cell culture was transfected with plasmids expressing gRNAs. Each experimental gRNA was transfected individually with two biological replicates, while control gRNAs were pooled equally and transfected together. 24 hours after transfection, the genomic DNA was extracted from transfected cells using the QuickExtract DNA Extraction Solution (Lucigen, QE09050). Targeted PCR using genomic DNA were performed to amplify gRNA-targeting regions, followed by index PCR to add adaptors for sequencing with iSeq 100 (∼10,000 reads per sample/replicate; Illumina). The analysis was done using customized scripts to only include frameshift indels^93^. Oligos used for targeted PCR are listed in Supplementary Table 7.

### Lentiviral and AAV vector construction, production, and QC

Viral vectors and plasmids were constructed as previously reported^24^. The backbone plasmid contains a human U6 promoter to express gRNAs and an EF1a promoter to express one or more labeling tags. The gRNA was designed as a dual-gRNA cassette, where two-gRNA constructs targeting the same gene (or two controls) are connected by a tRNA linker, adopted from a previous design^94^. The vectors were cloned individually and confirmed by Sanger sequencing. Lentiviral production and titration were performed by the Viral Core Facility at Sanford Burnham Prebys. AAV production, titration, gRNA distribution and related quality control assessments were performed as previously described^24^. Oligos used for gRNA amplicon targeted PCR are listed in Supplementary Table 7.

### Perturb-CLEAR labeling design

Multiple tags (tdTomato, DsRed, smFP.Myc, smFP.HA, smFP.V5, smFP.FLAG) were screened for labeling optimization^95,96^. Each tag was conjugated with a membrane-tethered motif to achieve plasma membrane localization^34,37^. The membrane-tethered tags were first screened *in vitro* and *in vivo* for their labeling performance and staining quality. tdTomato, smFP.Myc, and smFP.HA were used in the morphological analysis experiment. The ten NDD genes were split into two subsets (subset1: *Grin2b*, *Adnp*, *Arid1b*, *Shank3*, *Med13L*; subset2: *Syngap1*, *Foxp1*, *Kmt5b*, *Pogz*, *Cux2*) to overcome the limitation of imaging channels. Specifically, *Grin2b* and *Cux2* were exchanged in batch BW106 and BW107 (Supplementary Table 8). Three membrane-tethered tags (tdTomato-m, HA-m, Myc-m) and their pairwise combinations (Myc-m-P2A-tdTomato-m, HA-m-P2A-tdTomato-m, Myc-m-P2A-HA-m), yielding six tag variants in total, were used to barcode five perturbations and one control (Extended Data Fig. 9D). For the control condition, three safe-targeting and three non-targeting control vectors were pooled and assigned to a single tag.

### *In utero* administration of viral vectors

AAV or lentivirus (0.5-1.5 µL per embryo) was administered *in utero* to the lateral ventricle at E13.5 or E15.5 as previously described^24^. AAV stock concentration is typically 1-4×10^12^ genome copies (GC) /mL, whereas lentiviral stock concentration is typically 2.7×10^6^-3×10^8^ FU/mL. Particularly, for single tag injections, around 3×10^6^ FU/mL is optimal to maintain sparse labeling, as higher titers result in dense labeling with overlapping dendritic arbors, complicating morphological tracing. For multiplex labeling (tag variants > 3), injection titer can be increased to around 7×10^7^ FU/mL as overlapping neurons may carry different tags and can be resolved through distinct imaging channels (as used for the Perturb-CLEAR NDD risk gene screen in this study).

### Tissue harvest, staining and imaging

*Tissue collection, clearing, and whole-mount staining*: Postnatal pups and adult mice were anesthetized and transcardially perfused with ice-cold PBS followed by ice-cold 4% paraformaldehyde in PBS. The brain samples were dissected and postfixed overnight in 4% paraformaldehyde in PBS at 4 °C with rocking. All following steps were performed at room temperature (if not otherwise specified). Fixed brains were then washed in PBS for 2 hours. The hemisphere sample for the light-sheet dataset was processed by Lifecanvas Technologies. Other brain samples were vibratome sliced into 1 mm sections, then cross-linked and delipidated using the Clear+ Passive Clearing Kit (Lifecanvas Technologies, #C-PCK-500-1.52) according to the manufacturer’s protocol^97^. After de-lipidation, brain sections were washed 3 times in PBS with 0.3% Triton X-100 (0.3% PBST) overnight at 37 °C with rocking. For staining, the samples were first incubated in blocking media (10% donkey serum, 5% bovine serum albumin in 0.3% PBST) overnight. The samples were then incubated in 1 μg/mL primary antibodies in the blocking media for 2 days, followed by washing in 0.3% PBST 5 times for a total of 1 day. Lastly, the samples were then incubated in 2 μg/mL secondary antibodies in blocking media for 2 days, followed by washing in 0.3% PBST 5 times for a total of 1 day. After staining, samples were incubated in EasyIndex (Lifecanvas Technologies, #EI-100-1.52) for 1-2 hours or until the samples were optically clear. Specifically for brain samples from Cas9 mice, the de-lipidated sections were photobleached for two days using a 48 W LED light source to remove endogenous fluorescent signals.

The primary antibodies and dilutions were: Rabbit anti-RFP (Rockland 600-401-379, 1:500), Goat anti-RFP (Rockland 200-101-379, 1:500), Rabbit anti-Myc tag (Sigma C3956, 1:500), Rabbit anti-HA tag (Cell Signaling C29F4, 1:500), Rat anti-HA tag (Roche 11867423001, 1:100-200), Rabbit anti-V5 tag (Cell Signaling D3H8Q, 1:500), Mouse anti-FLAG tag (Sigma F1804, 1:1000), Rabbit anti-GFP (Invitrogen A-11122, 1:500).

Whole-mount imaging and image processing: The light-sheet data were generated by LifeCanvas Technology LLC using a SmartSPIM light-sheet microscope using 4× and 9× objectives. For whole-mount imaging of 1 mm brain sections, a Nikon AX Confocal Microscope and an Oxford Instruments Andor BC43 Microscope were used for 1 mm thick brain section imaging. For each brain section, an overview was first taken using a 10× air objective, followed by zoom-in imaging of the somatosensory cortex (visually identified by the high autofluorescence from the barrel cortex structure) using 20× air or water objective, with an N.A. value > 0.8.

### Neuron morphology 3D reconstruction and quantification

Soma positions of labeled neurons were annotated using the Filaments tool in Imaris 10.2.0 (Oxford Instruments). The boundary between layers 4 and 5 was defined by the strong autofluorescence of the barrel cortex (Extended Data Fig. 4A), and neurons with somata located below the lower edge of this structure were excluded. Inhibitory neurons or spiny stellate neurons were excluded based on their stereotypic morphology. The annotated soma positions were then used to segment 3D image volumes of individual neurons (approximately 700 × 700 × 450 μm^3^) with a customized script (See Code Availability). Images were removed during quality control if the signal-to-noise ratio was too low for reliable tracing (usually < 1.5) or if dendritic arbors were visibly truncated. Morphological reconstructions were subsequently generated from these single-neuron image volumes in Imaris 10.2.0 or neuTube 1.0. To ensure high data quality, reconstructions were generated first semi-manually by a trained reconstruction team, then independently proofread through visual inspection and manually corrected by at least two experienced analysts. A final quality-control step was performed using a customized script (See Code Availability) to identify and correct reconstruction artifacts, including over-branched nodes (> 2 daughter branches), spurious short branches (< 1 μm), and disconnected dendritic compartments. Specifically, whole neuron projection reconstruction (Extended Data Fig. 3E) was generated directly in Imaris without additional segmentation.

### Morphometrics data processing

#### Metric selection

A complete set of 170 metrics was designed to cover various aspects of neuronal morphology, including size, complexity, branching angles, structural heterogeneity, etc. (Supplementary Table 1). Derived summary-statistic variants (minimum, median, maximum, and standard deviation) of other existing metrics were excluded prior to analysis. The raw metrics matrix was imputed for missing values using k-nearest neighbor and scaled by z-score within each metric. Pairwise Pearson correlations were then computed from the remaining 98 metrics, which were used for hierarchical clustering. Modules were defined using an adaptive tree-cutting procedure using cutreeDynamic from the dynamicTreeCut package (1.63-1)^98^. Statistical significance of module separation was assessed via permutation testing. A reduced metric panel (MetricSet1, n = 36) was then selected for phenotypic tests. MetricSet1 was chosen to: (1) include at least one metric from each module, (2) minimize redundant readouts within modules, and (3) prioritize interpretable, commonly used metrics. Specifically, some metrics were measured separately for each dendritic compartment (Basal, Tuft, Oblique, or Global) to enable dendritic compartment-resolved quantification.

#### Raw data processing and clustering

Morphometric measurements and cell-level metadata were imported into R (v4.0.3)^99^. Missing values in the morphometric matrix were imputed by replacing each NA with the mean value of the corresponding metric across all cells. The resulting matrix was standardized using the scale function, and unsupervised clustering was performed with kmeans. Principal component analysis (PCA) was carried out on the scaled data using prcomp, and principal component scores were extracted for downstream analysis and visualization. To assign cell type labels, morphological metrics (MetricSet1, n = 36) were extracted from L2/3 and L4/5 neuron traces of Scala *et al.* 2021^20^, which were then imported into R for mapping analysis. A support vector machine (SVM) classifier was trained using the svm function from the e1071 package (v1.7-4)^100^, with scaled morphological metrics as predictors and RNA.type as the response variable. The trained model was then applied to the cells in this study, and cells were classified as L2/3 IT or L4/5 IT based on the predicted label.

#### Morphometrics heatmap visualization

Heatmaps were generated to visualize morphological metrics across neurons and subtypes. Neurons (columns) were ordered by their subtype assignment, with subtype levels ranked by their mean Tuft branch numbers (descending). Within each subtype, neurons were further sorted according to the mean standardized signal across subtype “signature” metric (defined as metrics whose maximal subtype-specific mean occurred in that subtype). Heatmap values were derived from the raw measurements (without imputation). Each morphometric (row) was standardized across neurons using z-score (excluding missing values), and originally missing values were retained as NA. Morphometric features were ordered hierarchically using pairwise correlation distance (1 − Pearson correlation). For visualization, standardized values were clipped to ±3 to reduce the influence of extreme outliers, and missing values (NA) were rendered in gray.

#### Statistics test across metrics

For differential testing across morphological metrics, 909 neurons carrying perturbations or control gRNAs were subsetted and, for each morphometric, a linear mixed-effects model was fit in R using lmer from the lmerTest package (v3.1-3)^101^. Each model included perturbation as a fixed effect and brain of origin as a random effect, and test statistics were extracted using summary. For a given morphometric, neurons with missing values (NA) in the raw (non-imputed) matrix were excluded from that analysis. Resulting *P*_val_ were adjusted for multiple testing using the false discovery rate (FDR) method implemented in p.adjust. The same modeling framework was also used for comparisons across ages.

#### NMF analysis

For non-negative matrix factorization (NMF), raw metrics were first scaled using scale, after which missing values were imputed by replacing each NA with the mean value of the corresponding metric. NMF was then performed on the morphometric matrix using nmf from the RcppML package (v0.5.6)^102^. Statistical testing of differences in NMF factor scores across perturbations was performed using the same linear mixed-effects modeling approach described above, substituting NMF factors for individual morphological metrics.

### Spine quantification

Perturb-CLEAR labeled brain samples were perfused and fixed as previously described, then vibratome sectioned into 100 μm slices. The slices were incubated in the blocking media (10% donkey serum, 5% bovine serum albumin in 0.3% PBST) overnight at 4 ℃. Next, the slices were incubated with primary antibodies (diluted in the blocking media) for 1-2 days at 4 ℃, followed by washing in 0.3% PBST five times. The slices were then incubated with secondary antibodies (diluted in the blocking buffer or 0.3% PBST) overnight at 4 ℃, followed by washing in 0.3% PBST. Lastly, the slices were mounted and sealed for imaging. Images of dendritic spines were taken using Nikon AX Confocal Microscope with a 60× oil objective (N.A. = 1.42). Neurons with somata localized within layer 2/3 and layer 4 were identified visually, and a 3D stacked image was taken for each neuron with the z step = 0.5 μm. Prior to spine counting, dendrite crops were made for each neuron focusing on tertiary or quaternary dendrites in the basal compartments, with the length > 25 μm for each crop. Spine number was manually counted using ImageJ v1.54r^103^ for each dendrite, and then length-weighted to obtain an average for each neuron. To test age-dependent changes of spine density within each cell type, separate one-way ANOVAs were performed for each cell type. Tukey’s post hoc multiple-comparisons test was applied to calculate *P*_adj_.

### Wild-type developmental transcriptomic data processing and analysis

#### Data Collection

Processed single-cell RNA sequencing (scRNA-seq) data were provided by the authors of Gao *et al.* 2025^59^. Despite potential region-specific differences, recent postnatal atlases indicate that somatosensory and visual cortex share broadly conserved maturation trajectories^60^, supporting our choice of using this as a developmental reference for somatosensory cortex. Analyses were restricted to 83,168 L2/3 IT neurons and 111,340 L4/5 IT neurons. Pseudobulk Differential Gene Expression Analysis: Differential gene expression analysis was performed across postnatal ages P10 (represented by combining P9-11 samples due to the limited sample number), P14, and P28 at the pseudobulk level using the AggregateExpression function from the Seurat R package (v5.3.1)^104^. Raw counts of cells were aggregated by biological sample and postnatal age. Genes were filtered by retaining genes with at least 10 counts in a minimum of two pseudobulk samples and differential gene expression analysis was performed using the DESeq2 (v1.42.1)^105^ package. The DESeq function was run using default settings, including median-of-ratios normalization for size factor estimation and parametric dispersion fitting.

#### Gene set enrichment analysis

Gene set enrichment analysis (GSEA) was performed using the gseGO function from the clusterProfiler R package (v4.10.1)^106^ using ranked gene lists derived from log_2_(Fold Changes). Gene Ontology (GO) terms with a minimum gene size of 10 and a maximum gene set size of 5,000 were included in the analysis.

#### Classification of temporal expression dynamics

Up-regulated DEGs in P14 vs. P9-11 comparison (*P*_adj_ < 0.05 and log_2_(Fold Change) > 0.3) were included for temporal analysis. Pseudobulk-level average normalized expression values were calculated for each biological sample across postnatal ages. Expression values were scaled relative to the P14 time point by subtracting the P14 expression value and dividing by the gene-specific standard deviation across samples. Based on log_2_(Fold Changes) between P14 and P28, genes were assigned to one of three temporal groups: Progressing (log_2_(Fold Change) > 0.3), Stabilizing (−0.3 < log_2_(Fold Change) < 0.3), or Transient (log_2_(Fold Change) < −0.3). Expression trends were summarized across samples by computing smoothed curves using locally weighted regression with a span parameter of 0.3.

#### Enrichment with selected gene sets

Risk genes associated with neurological disorders were collected from large-scale exome studies^61–64^. Human genes were converted to their corresponding mouse homologs prior to analysis based on the Ensembl BioMart homolog database^107^. Enrichment of risk genes with Progressing, Stabilizing, and Transient gene signatures was assessed using one-sided Fisher’s exact test, followed by multiple testing correction. Synapse-related gene set enrichment analysis was performed using the SynGO web-based platform^108^. Mouse genes were converted to their corresponding human homologs prior to analysis.

### *In vivo* Perturb-seq experiment

#### Nuclei isolation

Animals were deeply anesthetized with isoflurane, and the somatosensory cortex was dissected on ice. Nuclei isolation medium consisted of 250 mM sucrose (Sigma #84097), 25 mM KCl (Thermo Fisher Scientific #AM9640G), 10 mM HEPES (Thermo Fisher Scientific #15630080), and 5 mM MgCl_2_ (Sigma #M1028). Tissue was transferred to Dounce grinders (Sigma D8938) containing 1 mL homogenization buffer composed of 1 mM dithiothreitol (Sigma #646563), 0.1% IGEPAL CA-630 (Sigma #I8896), 1% BSA (Miltenyi #130-091-376), 10 mg/mL Kollidon VA64 (Sigma #190845), and 0.6 U/μL NxGen RNase Inhibitor (Biosearch Technologies #30281-1) in the nuclei isolation medium. Samples were dounced and homogenized, then incubated on ice for 3 min after addition of 1 mL fresh cold homogenization buffer. Approximately 2 mL homogenate was filtered through a 70 μm strainer (Greiner Bio-One #542170) into 4 mL cold wash buffer containing 1 mM dithiothreitol, 0.1% IGEPAL CA-630, 1% BSA, and 0.6 U/μL NxGen RNase Inhibitor in nuclei isolation medium. The suspension was centrifuged at 200×g for 5 minutes, the supernatant removed, and the pellet resuspended in 0.5 mL cold wash buffer supplemented with Alexa Fluor 647 rabbit anti-NeuN antibody (1:1000; Abcam ab190565). After 15 min on ice, nuclei were pelleted again at 100×g for 8 minutes, resuspended in 0.5 mL cold wash buffer containing 1 μg/mL DAPI (Thermo Fisher Scientific #D1306), and filtered into 5 mL test tubes (Corning #352235). GFP^+^, DAPI^+^, and NeuN^+^ nuclei were isolated by FANS using 100 μm sorting chips and collected into 0.5 mL cold wash buffer. Immediately after sorting, nuclei were centrifuged at 500×g for 8 minutes to remove excess supernatant before loading onto the 10x Chromium platform (10x Genomics).

#### Library construction and sequencing

Each channel for 10x Genomics snRNA-seq library was prepared by combining the FACS sorted nuclei from 1-2 animals. The nuclei isolation and FACS purification were performed within 2-3 hours under cold conditions to minimize nuclei damage. The batch1 library was generated using the Chromium Next GEM Single Cell 5’ Reagent Kits v2 with Feature Barcode Technology (10x Genomics). The batch2, 3, and 4 libraries were generated using the Chromium GEM-X Single Cell 5’ Reagent Kits v3 with Feature Barcode Technology (10x Genomics), according to the manufacturer’s protocol. Gene-expression and CRISPR gRNA libraries were pooled at a 10:1 ratio and sequenced either on an Illumina NextSeq 1000/2000 using XLEAP-SBS 100-cycle chemistry or by esBiolab paired-end 168-cycle sequencing, to a depth of > 22,000 reads per cell. Read lengths were 26 bases for Read 1 and 85 bases for Read 2 for Next GEM libraries, and 28 bases for Read 1 and 90 bases for Read 2 for GEM-X libraries. Index read lengths were 10 bases for i5 and i7 for Next GEM and GEM-X.

### Perturb-seq data processing and analysis

#### Data processing

Single-cell RNA-seq data were processed as previously described^24^. BCL files from transcriptome and gRNA feature-barcoding libraries were converted to FASTQ format using Illumina bcl2fastq with default parameters. Transcriptome and gRNA reads were aligned to the mouse reference genome mm10 (GENCODE vM23/Ensembl 98), and gene- and gRNA-expression count matrices were generated using outputs from 10x Genomics Cell Ranger v8.0.0^109^. Filtered count matrices were imported into R (v4.0.3) using Read10X from Seurat (v4.0.0)^104^ and used to create Seurat objects, excluding cells with < 250 detected genes. Data were log-normalized, and the 2,000 most variable features were identified using FindVariableFeatures. The integrated dataset was then scaled with ScaleData. Dimensionality reduction was performed using gbm.sc from scGBM (v0.1.0)^110^ with subset = 50000 and M = 25. Clustering was performed on the resulting scGBM reduction using FindNeighbors and FindClusters in Seurat, and UMAP embeddings were generated with RunUMAP. To identify putative doublets, cxds, bcds, and cxds_bcds_hybrid from the scds (v1.6.0)^111^ package were run separately on each 10X channel. The dataset was then uploaded to MapMyCells^112^ to obtain initial cell type annotations. Clusters were assigned cell type identities based on these labels, canonical marker gene expression, and quality-control metrics. Clusters identified as putative doublets or as having elevated mitochondrial content (identified by plotting percent mitochondrial RNA with a FeaturePlot and ViolinPlot and locating clusters with relatively high percent mitochondria) were annotated accordingly. For downstream analyses, cells assigned to doublet or high-mitochondrial clusters were excluded, as well as cells assigned to multiple perturbations, cells with a scds score > 1.2, cells with > 30,000 UMIs, and cells with > 2% mitochondrial UMIs.

#### gRNA assignment

Initial perturbation assignments were extracted from Cell Ranger outputs. For each paired set of guides, a combined guide label was generated, and cells assigned as singlets and assigned to either guide in the pair were reassigned to the corresponding combined guide. Cells initially assigned as doublets but assigned specifically to both guides within the same pair were likewise reclassified as singlets and assigned to the combined guide. Reliable perturbation assignment was supported by three observations: 1. the two gRNAs encoded on the same vector were frequently co-detected across cells, as shown by a high Pearson correlation between their Cell Ranger UMI counts; 2. among cells with two assigned gRNAs (based on Cell Ranger assignment), the assigned guides often corresponded to the matched pair from the same vector (36.4% of cells with two assigned gRNAs were assigned to the paired gRNAs, whereas if this was by chance it would be only 1.93%); 3. cells assigned to each perturbation were enriched for the expected target-site indels (Extended Data Fig. 12G-H). Indel analysis was performed as reported previously^31^.

#### Differential gene expression

Differential expression analysis was run in cell types with > 500 cells and for perturbations with > 10 cells within a given cell type, restricting analysis to genes expressed in at least 1% of cells. The safe-targeting control 2 (control-ST2) was set as the reference perturbation. A design matrix was constructed from the metadata, including perturbation and batch terms. Count data were written to disk as HDF5 files using writeHDF5Array from HDF5Array (v1.34.0)^113^ and reloaded as HDF5Array objects. These HDF5Array objects, together with the design matrix, were supplied to glm_gp from glmGamPoi (v1.18.0)^114^ with verbose = TRUE, on_disk = TRUE, and subsample = TRUE, using otherwise default settings to fit a negative binomial model. For each perturbation, log_2_(Fold Changes) and *P*_adj_ were obtained using test_de with pval_adjust_method = “BH” and compute_lfc_se = FALSE.

### Foxp1 DE gene log fold change (logFC) comparison

Data from Ortiz *et al.* 2025^72^ were downloaded and loaded into Seurat (v4.0.0)^104^. Clustering was performed as described for the Perturb-seq data. Cell type annotations were obtained directly from MapMyCells^112^ and used for downstream analyses. Within each cell type, differential expression was assessed from pseudobulk profiles generated with AggregateExpression using slot = “counts”. Genes detected at > 10 UMIs in fewer than three samples were excluded, after which normalization factors were computed with calcNormFactors from edgeR (v3.32.1)^115^. Additional gene filtering was performed with filterByExpr, dispersion was estimated with estimateDisp, model fitting was carried out with glmFit, and differential expression testing was performed with glmLRT. Multiple-testing correction was applied using p.adjust (method = “fdr”), and empirical *P*_val_ were computed with fdrtool (v1.2.16)^116^. Nominally significant DE genes were defined using an uncorrected threshold (*P*_val_ < 0.05) and FDR-supported DE genes were defined using an adjusted threshold (*P*_adj_ < 0.05). For visualization, log_2_(Fold Changes) of all shared-nominal genes from the two datasets were plotted. All statistical analyses were restricted to the shared FDR-supported gene set, including Pearson correlation (two-sided test) and sign concordance (fraction of genes with matching log_2_(Fold Changes) sign across datasets). A fitted regression line and its 95% confidence band, estimated with predict.lm, were overlaid on the scatter plot.

### Cross-modality joint analysis

#### TRADE analysis and correlation

TRADE analysis was performed on both morphometric and transcriptomic datasets. The TRADE function from the TRADE package (v0.1.0)^117^ was applied to a data frame containing *P*_val_, effect sizes, and standard errors, using mode = “univariate”, and the transcriptome_wide_impact value was extracted from the distribution_summary output. For transcriptomic TRADE analysis, differential expression analysis was rerun with compute_lfc_se = TRUE to recover standard errors before running TRADE. Pearson correlation was then computed across all perturbation and cell type combinations.

#### Energy distance calculation and correlation

For energy-distance analysis of the RNA-seq data, batch-corrected embeddings were first generated with RunHarmony from the Harmony package (v1.0)^118^. For the morphometric data, batch effects attributable to brain of origin were corrected using ComBat from the sva package (v3.38.0)^119^. In each case, energy distances were computed with edist from the scperturbR package (v0.1.0)^120^. For the morphometric dataset, we used a modified implementation of edist that accepts matrices rather than Seurat objects. The output in each case was a matrix of pairwise energy distances among perturbations and controls.

## Data availability

The gRNA sequences, raw metrics of morphology data, morphometric analysis results, and transcriptomic analysis results are included as Supplementary Tables 2-14. Single-nucleus RNA-seq data are available through the Gene Expression Omnibus and UCSC cell browser. The 3D reconstruction traces of neuronal morphology (.swc files) are available through Brain Imaging Library.

## Code availability

Codes used for analysis for this study are available at Github (https://github.com/jinlabneurogenomics/Perturb-CLEAR).

## Acknowledgements

We thank Jorryn Wu, Chloe Samouhi, Andrea Balcan, Qiucun She, Siena Parr, Allyna Huang, Adeline Sun, Lori Demirjian, Hayley Prinstein, Fiona Gan, Daniel Nguyen, and Tianna Nguyen for their help with neuronal tracing; Jakub Ziak, Alex Kolodkin, Li Ye, and Ardem Patapoutian for their advice on the manuscript; Scripps Research Department of Animal Resources, DNC Microscopy Core and Shakib Omari, Genomics Core, and Flow Cytometry Core for assistance; Zhengyuan Pang for the help with data visualization; and all members of the Jin lab for their support. The lentiviral production in this work was by the Sanford Burnham Prebys Functional Genomics Core and supported by the Shared Instrumentation Grant S10-OD036254 and the NCI Cancer Center Support Grant P30-CA030199. B.W. was supported by the Mark Pearson Endowed Fellowship. J.L. was supported by CIRM Training Fellowship (EDU4-12811) and Dorris Scholar Award. X.Z. was supported by Dorris Scholar Award and the Frank J. Dixon Fellowship. X.W.Y was supported by the National Institutes of Health BRAIN Initiative grants (U01MH117079, RF1MH128888). J.Z.L. was supported by the Stanley Center for Psychiatric Research at the Broad Institute of MIT and Harvard. J.Z.L and X.J. were supported by the National Institutes of Health (R01MH137042) and are part of the SSPsyGene Consortium. X.J. and this work were supported by HHMI, Simons Foundation Autism Research Initiative Sex Differences Collaboration (SFARI #736613), CIRM (DISC0-14424, ReMIND-L DISC4-16295), National Institutes of Health (R01HG012819, R01AT013748), the Mathers Foundation, the One Mind Foundation, the Conrad Prebys Foundation, the Pew Charitable Trust, the McKnight Endowment, the Astera Institute, Jed McCaleb, and James Fickel.

## Author contributions

B.W. and X.J. conceived the project and designed the experiments with input from all authors. B.W., N.H., and G.S.C. constructed and produced AAV vectors; B.W. performed in vivo experiments with the help of X.J.; B.W., J.L., M.A.A., C.S.P., I.M., S.P., A.C., N.B., P.D., and T.N. performed tissue collection, staining, imaging, and raw data collection; B.W. and S.K.S. performed morphology analysis under the supervision of X.W.Y., J.Z.L., and X.J.; B.W. and S.K. performed developmental transcriptomic analysis under the supervision of X.J.; B.W. performed snRNA-seq experiments with the help from J.L. and X.Z.; B.W., S.K.S., and J.L. performed Perturb-seq analysis under the supervision of J.Z.L. and X.J.; B.W. and X.J. drafted the manuscript with input from all authors.

---

**Extended Data Fig. 1. *In vitro* and *in vivo* screen of labeling reporters with consistent and high-performance cytoarchitectural readouts.**

A. Immunostaining showing membrane-tethered reporters labeling in transfected HEK293T cells. tdTomato-m: membrane-tethered tdTomato, DsRed-m: membrane-tethered DsRed. Myc-m: membrane-tethered smFP.Myc, HA-m: membrane-tethered smFP.HA, V5-m: membrane-tethered smFP.V5, FLAG-m: membrane-tethered smFP.FLAG. tdTomato-m, Myc-m, HA-m, and V5-m were used for *in vivo* test. Scale bar: 20 μm.

B. Immunostaining showing membrane-tethered reporters labeling in transduced neurons *in vivo*. Scale bar: 50 μm.

C. Correlation plot showing high consistency between the SD-scaled means of MetricSet1 derived from Perturb-CLEAR-labeled neurons using different tags. A fitted linear regression line is shown with 95% confidence band. Dashed line indicates y = x. tdTomato-m and HA-m-labeled wild-type P10 L4/5 IT neurons are used as examples, tdTomato-m: n = 7 from N = 3 brains; HA-m: n = 9 from N = 4 brains.

**Extended Data Fig. 2. Viral titer optimization allows appropriate labeling density for dendritic mapping and reliable perturbation identity assignment.**

A. Immunostaining showing the labeling density in somatosensory cortex across titers (2.7×10^6^– 1.5×10^8^ FU/mL). Maximum intensity projections from 400 μm thick slab from whole-mount images. Scale bar: 200 μm.

B. Correlation between injection titers and labeling density of Perturb-CLEAR brain samples. Titer is plotted on a log10 scale, whereas density is shown on the original scale. The interquartile range (IQR) with median (center line) is shown, with whiskers extending to 1.5× IQR; overlaid jittered points indicate individual brains. A fitted linear regression line (solid) is shown with 95% confidence band. n > 3 brains for each condition. FU: functional unit.

C. Immunostaining of somatosensory cortex from a multiplex labeled brain sample (mixed lentiviral vectors with tdTomato-m, V5-m, and Myc-m, 3.0×10^8^ FU/mL). Maximum intensity projections are from 100 μm brain sections. Scale bar: 100 μm.

D. Quantification showing low double-positive rate despite a high injection titer (3.0×10^8^ FU/mL). N = 3 brains.

**Extended Data Fig. 3. Perturb-CLEAR allows brain-wide cytoarchitecture mapping across cell types, brain regions, and scales.**

A. Overview of a Perturb-CLEAR brain sample imaged by light-sheet microscopy.

B. Zoom-in maximum projections from Fig. 1B showing laminar distribution of deep layer cortical pyramidal neurons. Scale bar: 100 μm.

C. Same as Extended Data Fig. 3B but from different regions of interest. CTX Inh, cortical inhibitory neurons; TH, thalamus; CB, cerebellum. Scale bar: 100 μm.

D. Zoom-in maximum projections show axon tracts (lower left) and dendritic spines (lower right). Scale bar: 20 μm.

E. Long-range axonal projection tracing of Perturb-CLEAR labeled L4 (intratelencephalic) and L5 (extratelencephalic) pyramidal neurons. Scale bar: 1 mm.

**Extended Data Fig. 4. Perturb-CLEAR dataset sample summary and raw traces.**

A. Representative immunostaining images show somatosensory cortex segmentation and cortical layer annotation based on barrel cortex autofluorescence (Green). Coarse dashed line: somatosensory cortex segmentation. Fine dashed line: boundary between layer 4 and layer 5. Scale bar: 1 mm.

B-C. UMAP embeddings and bar plot showing sample distribution across batches (animal litters).

D. Representative traces showing consistent identification of morphological subtypes across batches.

E. Bar plot showing proportion of morphological subtypes across batches.

**Extended Data Fig. 5. Morphometric correlation analysis identifies metric modules.**

Heatmap showing the Pearson *R* derived from pairwise correlation between metrics from the Perturb-CLEAR dataset (n = 1053 neurons, n = 98 metrics). “Selected” annotation bar indicates metrics that are included in the MetricSet1 for statistical test.

**Extended Data Fig. 6. Quality control and sample distribution of the developmental morphology analysis.**

A. Bar plot showing the brain distribution of Perturb-CLEAR neurons used in the developmental analysis (Fig. 2B).

B. Bar plot showing the percentage of cells assigned to each inferred cell type for the developmental analysis (n = 144 neurons) (related to Fig. 2B).

C. Dot plot showing shared and distinct morphological differences comparing P14 to P10 and P28, across wild-type L2/3 IT and L4/5 IT neurons. Different gray shadings indicate categories of morphometric features (size, complexity, and angle). P10: n = 21 L2/3 IT neurons from 4 brains, n = 17 L4/5 IT neurons from 4 brains. P14: n = 53 L2/3 IT neurons from 3 brains, n = 11 L4/5 IT neurons from 3 brains. P28: n = 28 L2/3 IT neurons from 4 brains, n = 14 L4/5 IT neurons from 3 brains.

**Extended Data Fig. 7. Dendritic compartment- and cell type-specific morphological development dynamics.**

A-C. Plots showing the developmental dynamics of representative metrics from Perturb-CLEAR neurons used in the developmental analysis (Fig. 2B and Extended Data Fig. 6C). Metrics were plotted over numeric postnatal ages, with conditions offset slightly for visualization purposes. Raincloud plots show half violins and half scattered points (individual neurons). Diamonds mark group means, and error bars represent t-based 95% confidence interval. *: *P*_adj_ < 0.05; ns: *P*_adj_ > 0.05.

**Extended Data Fig. 8. Characterize gene signature using wild-type developmental mouse brain RNA data.**

A. Histogram showing the distribution of all genes (upregulated genes comparing P14 to P9-11) by log_2_(Fold Change) comparing P28 to P14. Dashed lines indicate selection criteria for each gene signature. *Progressing*: log_2_(Fold Change) > 0.3, *Transient*: log_2_(Fold Change) < −0.3, *Stabilizing*: −0.3 < log_2_(Fold Change) < 0.3.

B. Developmental expression trajectories of genes within the *Progressing* (L2/3 IT: n = 1292; L4/5 IT: n = 1240) and the *Transient* signatures (L2/3 IT: n = 637; L4/5 IT: n = 570). Scaled expression level relative to P14 time point is shown.

**Extended Data Fig. 9. Quality control and sample distribution of the morphological screen.**

A. Dot plots showing developmental expression trajectories of selected genes across cell types.

B. Heatmap showing the on-target frameshift indel rate of selected gRNA in transfected cell culture. Each gRNA was measured with two biological replicates (Rep.1 and 2). The control condition was transfected by a mixture of control gRNAs including safe-targeting and non-targeting. See Methods for details.

C. Histogram showing the gRNA distribution of the pooled lentiviral vectors with the kernel density estimation (KDE), sampled from purified lentiviral supernatant.

D. Representative immunostaining images of neurons expressing different tags or tag combinations. Scale bar: 50 μm.

E. Bar plot showing the distribution of neurons from morphological screen (n = 909) across batches (animal litters).

F. Bar plot showing the percentage of cells assigned to each inferred cell type across perturbations and control.

**Extended Data Fig. 10. Phenotype summary of cell type- and dendritic compartment-specific morphological changes introduced by NDD risk genes.**

A. Dot plot showing that NDD risk gene perturbations introduce specific morphological changes in L2/3 IT neurons (n = 804). Different gray shadings indicate categories of morphometric features (size, complexity, and angle). Sample details were included in Supplementary Table 8.

B. Dot plot showing that NDD risk gene perturbations introduce specific morphological changes in L4/5 IT neurons (n = 105). Different gray shadings indicate categories of morphometric features (size, complexity, and angle). Sample details were included in Supplementary Table 8.

**Extended Data Fig. 11. Phenotype summary of *Syngap1*- and *Adnp*-specific morphological changes.**

A. Box plot showing morphological changes comparing *Syngap1*-perturbed and control neurons across metrics. The interquartile range (IQR) with median (center line) is shown, with whiskers extending to 1.5× IQR; overlaid jittered points indicate individual neurons. *Syngap1*-perturbed: n = 68 neurons from 9 brains, control: n = 135 neurons from 19 brains. *: *P*_adj_ < 0.05.

B. Representative traces of *Syngap1*-perturbed and control L4/5 IT neurons.

C. Box plot showing basal height is not changed in *Syngap1*-perturbed L4/5 IT neurons. The interquartile range (IQR) with median (center line) is shown, with whiskers extending to 1.5× IQR; overlaid jittered points indicate individual neurons. *Syngap1*-perturbed: n = 22 neurons from 8 brains, control: n = 27 neurons from 10 brains. ns: *P*_adj_ > 0.05.

D. Heatmap of the weight matrix of NMF modules showing various enrichment of metrics onto each module in L2/3 IT neurons.

E. Dot plot of NMF analysis showing perturbation-specific module enrichment in L2/3 IT neurons.

F. Representative traces of *Adnp*-perturbed L4/5 IT neurons from the validation dataset.

G. Box plot showing morphological changes comparing *Adnp*-perturbed and control L4/5 IT neurons from the validation dataset. across metrics. The interquartile range (IQR) with median (center line) is shown, with whiskers extending to 1.5× IQR; overlaid jittered points indicate individual neurons. *: *P*_adj_ < 0.05; ns: *P*_adj_ > 0.05.

**Extended Data Fig. 12. Quality control and sample distribution of *in vivo* Perturb-seq.**

A. Histogram showing the gRNA distribution of the pooled AAV with the kernel density estimation (KDE) line.

B. Immunostaining showing AAV-transduced neurons distribution throughout the neocortex post in-utero injection. Scale bar: 1 mm.

C. Immunostaining showing a FANS-enriched GFP^+^ and NeuN^+^ neuronal nucleus. Scale bar: 10 μm.

D. Representative FANS profiles showing the gating strategies and an expected positive cell percentage for sorting.

E. UMAP embeddings showing the cell type clustering of all nuclei (n = 161,051 nuclei in total, n = 150,531 after removing clusters with high mitochondrial UMIs (high mito) and doublets).

F. UMAP embeddings showing data distribution across channels. Upper: all nuclei, bottom: only showing nuclei assigned with single perturbations.

G. Bar plot showing percentage of perturbation assignment across cell types.

H. UMAP embeddings showing the distribution of gRNA-assigned cells in clustering.

I. Heatmap showing pairwise detection correlation across gRNA species.

J. Bar plot showing transcript indels enrichment in assigned nuclei compared to others. Only gRNAs targeting transcript regions with sufficient read counts in the 5’ RNA-seq data are shown.

K. Violin plot showing the distribution of total gene number per cell (nGene), total transcript number per cell (nUMI), percentage of mitochondrial UMIs (mito%), and total transcript number of gRNA per cell, across cell types.

**Extended Data Fig. 13. Cell type composition analysis of *in vivo* Perturb-seq.**

A. Bar plot showing cell type percentage across perturbations.

B. Dot plot showing the cell number retrieved for each cell type post perturbation assignment.

**Extended Data Fig. 14. Phenotype summary of *in vivo* Perturb-seq.**

A. Volcano plots showing diverse DEGs introduced by *Foxp1* perturbation across cell type.

B. Correlation of the DEG log_2_(Fold Change) derived from *Foxp1* perturbed L5 PT neurons from this Perturb-seq data and those from *Foxp1* cKO L5 PT neuron from a public dataset^72^. All shared-nominal DEGs are shown. Only shared-FDR significant DEGs were used for correlation calculation and sign concordance. A fitted linear regression line is shown with 95% confidence band. Dashed lines indicate x = 0, y = 0, and y = x.

C. Bar plot of *Grin2b* perturbation DEG number across cell types shows cell type-specific impact on L6 CT.

D. Volcano plot showing DEGs introduced by *Adnp* perturbation in L2/3 IT neurons.

**Extended Data Fig. 15. Convergent and divergent phenotypic effects by NDD gene perturbation.**

A. Heatmap of energy distance (eDist) showing diverse phenotypic effects across perturbations in the morphological modality (black) or transcriptomic modality (red).

B. Volcano plot showing DEGs introduced by *Arid1b* perturbation in L4/5 IT neurons.

## Supplementary materials

Supplementary Table 1. Morphological metrics list

Supplementary Table 2. Developmental morphology sample summary, raw metrics, and test results

Supplementary Table 3. Spine analysis raw metrics and test results

Supplementary Table 4. Developmental transcriptome dataset DE genes

Supplementary Table 5. Gene list for each signature

Supplementary Table 6. Enrichment results (GSEA-GO, fisher exact test)

Supplementary Table 7. Plasmid, gRNA, oligonucleotides, and viral vectors

Supplementary Table 8. NDD screen and *Adnp* validation sample summary

Supplementary Table 9. NDD screen raw metrics and test results

Supplementary Table 10. *Adnp* validation raw metrics and test results

Supplementary Table 11. Perturb-seq sample summary and metrics

Supplementary Table 12. Perturb-seq DE genes

Supplementary Table 13. *Foxp1* transgenic model DE genes and analysis

Supplementary Table 14. Joint analysis metrics

## Reference

1 Zeng, H. What is a cell type and how to define it? Cell 185, 2739–2755 (2022). 10.1016/j.cell.2022.06.031

2 Luo, L. Architectures of neuronal circuits. Science 373, eabg7285 (2021). 10.1126/science.abg7285

3 Petreanu, L., Mao, T., Sternson, S. M. & Svoboda, K. The subcellular organization of neocortical excitatory connections. Nature 457, 1142–1145 (2009). 10.1038/nature07709

4 Rah, J. C. et al. Thalamocortical input onto layer 5 pyramidal neurons measured using quantitative large-scale array tomography. Front Neural Circuits 7, 177 (2013). 10.3389/fncir.2013.00177

5 Balcioglu, A. et al. Mapping thalamic innervation to individual L2/3 pyramidal neurons and modeling their ‘readout’ of visual input. Nat Neurosci 26, 470–480 (2023). 10.1038/s41593-022-01253-9

6 London, M. & Hausser, M. Dendritic computation. Annu Rev Neurosci 28, 503–532 (2005). 10.1146/annurev.neuro.28.061604.135703

7 Spruston, N. Pyramidal neurons: dendritic structure and synaptic integration. Nat Rev Neurosci 9, 206–221 (2008). 10.1038/nrn2286

8 Cline, H. T. Dendritic arbor development and synaptogenesis. Current Opinion in Neurobiology 11, 118–126 (2001). 10.1016/S0959-4388(00)00182-3

9 Flavell, S. W. & Greenberg, M. E. Signaling mechanisms linking neuronal activity to gene expression and plasticity of the nervous system. Annu Rev Neurosci 31, 563–590 (2008). 10.1146/annurev.neuro.31.060407.125631

10 Holtmaat, A. & Svoboda, K. Experience-dependent structural synaptic plasticity in the mammalian brain. Nat Rev Neurosci 10, 647–658 (2009). 10.1038/nrn2699

11 Jan, Y. N. & Jan, L. Y. Branching out: mechanisms of dendritic arborization. Nat Rev Neurosci 11, 316–328 (2010). 10.1038/nrn2836

12 Riccomagno, M. M. & Kolodkin, A. L. Sculpting neural circuits by axon and dendrite pruning. Annu Rev Cell Dev Biol 31, 779–805 (2015). 10.1146/annurev-cellbio-100913-013038

13 Lefebvre, J. L., Sanes, J. R. & Kay, J. N. Development of dendritic form and function. Annu Rev Cell Dev Biol 31, 741–777 (2015). 10.1146/annurev-cellbio-100913-013020

14 Kolodkin, A. L. & Tessier-Lavigne, M. Mechanisms and molecules of neuronal wiring: a primer. Cold Spring Harb Perspect Biol 3 (2011). 10.1101/cshperspect.a001727

15 Santiago, C. & Bashaw, G. J. Transcription factors and effectors that regulate neuronal morphology. Development 141, 4667–4680 (2014). 10.1242/dev.110817

16 Ouzounidis, V. R., Prevo, B. & Cheerambathur, D. K. Sculpting the dendritic landscape: Actin, microtubules, and the art of arborization. Curr Opin Cell Biol 84, 102214 (2023). 10.1016/j.ceb.2023.102214

17 Dong, X., Shen, K. & Bulow, H. E. Intrinsic and extrinsic mechanisms of dendritic morphogenesis. Annu Rev Physiol 77, 271–300 (2015). 10.1146/annurev-physiol-021014-071746

18 Cadwell, C. R. et al. Electrophysiological, transcriptomic and morphologic profiling of single neurons using Patch-seq. Nat Biotechnol 34, 199–203 (2016). 10.1038/nbt.3445

19 Gouwens, N. W. et al. Integrated Morphoelectric and Transcriptomic Classification of Cortical GABAergic Cells. Cell 183, 935–953 e919 (2020). 10.1016/j.cell.2020.09.057

20 Scala, F. et al. Phenotypic variation of transcriptomic cell types in mouse motor cortex. Nature 598, 144–150 (2021). 10.1038/s41586-020-2907-3

21 Peng, H. et al. Morphological diversity of single neurons in molecularly defined cell types. Nature 598, 174–181 (2021). 10.1038/s41586-021-03941-1

22 Berg, J. et al. Human neocortical expansion involves glutamatergic neuron diversification. Nature 598, 151–158 (2021). 10.1038/s41586-021-03813-8

23 Willsey, H. R., Willsey, A. J., Wang, B. & State, M. W. Genomics, convergent neuroscience and progress in understanding autism spectrum disorder. Nat Rev Neurosci 23, 323–341 (2022). 10.1038/s41583-022-00576-7

24 Zheng, X., Li, J. & Jin, X. Functional Neurogenomics to Dissect Disease Mechanisms Across Models. Annu Rev Genomics Hum Genet 26, 189–216 (2025). 10.1146/annurev-genom-120823-125811

25 Robertson, C. E. & Baron-Cohen, S. Sensory perception in autism. Nat Rev Neurosci 18, 671–684 (2017). 10.1038/nrn.2017.112

26 Marco, E. J., Hinkley, L. B. N., Hill, S. S. & Nagarajan, S. S. Sensory Processing in Autism: A Review of Neurophysiologic Findings. Pediatric Research 69, 48–54 (2011). 10.1203/PDR.0b013e3182130c54

27 Petersen, C. C. H. The Functional Organization of the Barrel Cortex. Neuron 56, 339–355 (2007). 10.1016/j.neuron.2007.09.017

28 Feldmeyer, D. Excitatory neuronal connectivity in the barrel cortex. Front Neuroanat 6, 24 (2012). 10.3389/fnana.2012.00024

29 Jin, X. et al. In vivo Perturb-Seq reveals neuronal and glial abnormalities associated with autism risk genes. Science 370 (2020). 10.1126/science.aaz6063

30 Santinha, A. J. et al. Transcriptional linkage analysis with in vivo AAV-Perturb-seq. Nature 622, 367–375 (2023). 10.1038/s41586-023-06570-y

31 Zheng, X. et al. Massively parallel in vivo Perturb-seq reveals cell-type-specific transcriptional networks in cortical development. Cell 187, 3236–3248 e3221 (2024). 10.1016/j.cell.2024.04.050

32 Stelzer, E. H. K. et al. Light sheet fluorescence microscopy. Nature Reviews Methods Primers 1 (2021). 10.1038/s43586-021-00069-4

33 Moffitt, J. R., Lundberg, E. & Heyn, H. The emerging landscape of spatial profiling technologies. Nat Rev Genet 23, 741–759 (2022). 10.1038/s41576-022-00515-3

34 Livet, J. et al. Transgenic strategies for combinatorial expression of fluorescent proteins in the nervous system. Nature 450, 56–62 (2007). 10.1038/nature06293

35 Gong, H. et al. Continuously tracing brain-wide long-distance axonal projections in mice at a one-micron voxel resolution. Neuroimage 74, 87–98 (2013). 10.1016/j.neuroimage.2013.02.005

36 Winnubst, J. et al. Reconstruction of 1,000 Projection Neurons Reveals New Cell Types and Organization of Long-Range Connectivity in the Mouse Brain. Cell 179, 268–281 e213 (2019). 10.1016/j.cell.2019.07.042

37 Veldman, M. B. et al. Brainwide Genetic Sparse Cell Labeling to Illuminate the Morphology of Neurons and Glia with Cre-Dependent MORF Mice. Neuron 108, 111–127 e116 (2020). 10.1016/j.neuron.2020.07.019

38 Gao, L. et al. Single-neuron analysis of dendrites and axons reveals the network organization in mouse prefrontal cortex. Nat Neurosci 26, 1111–1126 (2023). 10.1038/s41593-023-01339-y

39 Qiu, S. et al. Whole-brain spatial organization of hippocampal single-neuron projectomes. Science 383, eadj9198 (2024). 10.1126/science.adj9198

40 Munoz-Castaneda, R. et al. Cellular anatomy of the mouse primary motor cortex. Nature 598, 159–166 (2021). 10.1038/s41586-021-03970-w

41 Park, C. S. et al. Dendritome mapping reveals the spatial organization of striatal neuron morphology. Nat Neurosci 28, 2628–2643 (2025). 10.1038/s41593-025-02085-z

42 Yao, Z. et al. A high-resolution transcriptomic and spatial atlas of cell types in the whole mouse brain. Nature 624, 317–332 (2023). 10.1038/s41586-023-06812-z

43 Major, G., Larkum, M. E. & Schiller, J. Active properties of neocortical pyramidal neuron dendrites. Annu Rev Neurosci 36, 1–24 (2013). 10.1146/annurev-neuro-062111-150343

44 Schuman, B., Dellal, S., Prönneke, A., Machold, R. & Rudy, B. Neocortical Layer 1: An Elegant Solution to Top-Down and Bottom-Up Integration. Annual Review of Neuroscience 44, 221–252 (2021). 10.1146/annurev-neuro-100520-012117

45 Ledderose, J. M. T. et al. Layer 1 of somatosensory cortex: an important site for input to a tiny cortical compartment. Cereb Cortex 33, 11354–11372 (2023). 10.1093/cercor/bhad371

46 Huang, S., Wu, S. J., Sansone, G., Ibrahim, L. A. & Fishell, G. Layer 1 neocortex: Gating and integrating multidimensional signals. Neuron 112, 184–200 (2024). 10.1016/j.neuron.2023.09.041

47 Gouwens, N. W. et al. Classification of electrophysiological and morphological neuron types in the mouse visual cortex. Nat Neurosci 22, 1182–1195 (2019). 10.1038/s41593-019-0417-0

48 Gao, L. et al. Single-neuron projectome of mouse prefrontal cortex. Nat Neurosci 25, 515–529 (2022). 10.1038/s41593-022-01041-5

49 Liu, L. et al. Connectivity of single neurons classifies cell subtypes in mouse brains. Nat Methods 22, 861–873 (2025). 10.1038/s41592-025-02621-6

50 Callaway, E. M. & Borrell, V. Developmental sculpting of dendritic morphology of layer 4 neurons in visual cortex: influence of retinal input. J Neurosci 31, 7456–7470 (2011). 10.1523/JNEUROSCI.5222-10.2011

51 Romand, S., Wang, Y., Toledo-Rodriguez, M. & Markram, H. Morphological development of thick-tufted layer v pyramidal cells in the rat somatosensory cortex. Front Neuroanat 5, 5 (2011). 10.3389/fnana.2011.00005

52 Kroon, T., van Hugte, E., van Linge, L., Mansvelder, H. D. & Meredith, R. M. Early postnatal development of pyramidal neurons across layers of the mouse medial prefrontal cortex. Sci Rep 9, 5037 (2019). 10.1038/s41598-019-41661-9

53 Richards, S. E. V. et al. Experience-Dependent Development of Dendritic Arbors in Mouse Visual Cortex. J Neurosci 40, 6536–6556 (2020). 10.1523/JNEUROSCI.2910-19.2020

54 Ciganok-Huckels, N. et al. Postnatal development of electrophysiological and morphological properties in layer 2/3 and layer 5 pyramidal neurons in the mouse primary visual cortex. Cereb Cortex 33, 5875–5884 (2023). 10.1093/cercor/bhac467

55 Schachtele, S. J., Losh, J., Dailey, M. E. & Green, S. H. Spine formation and maturation in the developing rat auditory cortex. J Comp Neurol 519, 3327–3345 (2011). 10.1002/cne.22728

56 Iascone, D. M. et al. Whole-Neuron Synaptic Mapping Reveals Spatially Precise Excitatory/Inhibitory Balance Limiting Dendritic and Somatic Spiking. Neuron 106, 566–578 e568 (2020). 10.1016/j.neuron.2020.02.015

57 Stern, E. A., Maravall, M. & Svoboda, K. Rapid Development and Plasticity of Layer 2/3 Maps in Rat Barrel Cortex In Vivo. Neuron 31, 305–315 (2001). 10.1016/S0896-6273(01)00360-9

58 Wen, J. A. & Barth, A. L. Input-specific critical periods for experience-dependent plasticity in layer 2/3 pyramidal neurons. J Neurosci 31, 4456–4465 (2011). 10.1523/JNEUROSCI.6042-10.2011

59 Gao, Y. et al. Continuous cell-type diversification in mouse visual cortex development. Nature 647, 127–142 (2025). 10.1038/s41586-025-09644-1

60 Butrus, S., Monday, H. R., Yoo, C. J., Feldman, D. E. & Shekhar, K. Molecular states underlying neuronal cell type development and plasticity in the postnatal whisker cortex. PLoS Biol 23, e3003176 (2025). 10.1371/journal.pbio.3003176

61 Kaplanis, J. et al. Evidence for 28 genetic disorders discovered by combining healthcare and research data. Nature 586, 757–762 (2020). 10.1038/s41586-020-2832-5

62 Satterstrom, F. K. et al. Large-Scale Exome Sequencing Study Implicates Both Developmental and Functional Changes in the Neurobiology of Autism. Cell 180, 568–584 e523 (2020). 10.1016/j.cell.2019.12.036

63 Fu, J. M. et al. Rare coding variation provides insight into the genetic architecture and phenotypic context of autism. Nat Genet 54, 1320–1331 (2022). 10.1038/s41588-022-01104-0

64 Singh, T. et al. Rare coding variants in ten genes confer substantial risk for schizophrenia. Nature 604, 509–516 (2022). 10.1038/s41586-022-04556-w

65 Hall, B. J., Ripley, B. & Ghosh, A. NR2B signaling regulates the development of synaptic AMPA receptor current. J Neurosci 27, 13446–13456 (2007). 10.1523/JNEUROSCI.3793-07.2007

66 Clement, J. P. et al. Pathogenic SYNGAP1 mutations impair cognitive development by disrupting maturation of dendritic spine synapses. Cell 151, 709–723 (2012). 10.1016/j.cell.2012.08.045

67 Llamosas, N. et al. SYNGAP1 Controls the Maturation of Dendrites, Synaptic Function, and Network Activity in Developing Human Neurons. J Neurosci 40, 7980–7994 (2020). 10.1523/JNEUROSCI.1367-20.2020

68 Li, X. et al. Foxp1 regulates cortical radial migration and neuronal morphogenesis in developing cerebral cortex. PLoS One 10, e0127671 (2015). 10.1371/journal.pone.0127671

69 Bennison, S. A. et al. The cytoplasmic localization of ADNP through 14-3-3 promotes sex-dependent neuronal morphogenesis, cortical connectivity, and calcium signaling. Mol Psychiatry 28, 1946–1959 (2023). 10.1038/s41380-022-01939-3

70 Michaelson, S. D. et al. SYNGAP1 heterozygosity disrupts sensory processing by reducing touch-related activity within somatosensory cortex circuits. Nat Neurosci 21, 1–13 (2018). 10.1038/s41593-018-0268-0

71 Vermaercke, B. et al. SYNGAP1 deficiency disrupts synaptic neoteny in xenotransplanted human cortical neurons in vivo. Neuron 112, 3058–3068 e3058 (2024). 10.1016/j.neuron.2024.07.007

72 Ortiz, A. et al. Cell-type-specific roles of FOXP1 in the excitatory neuronal lineage during early neocortical murine development. Cell Rep 44, 115384 (2025). 10.1016/j.celrep.2025.115384

73 Bacon, C. et al. Brain-specific Foxp1 deletion impairs neuronal development and causes autistic-like behaviour. Mol Psychiatry 20, 632–639 (2015). 10.1038/mp.2014.116

74 Mao, L., Takamiya, K., Thomas, G., Lin, D. T. & Huganir, R. L. GRIP1 and 2 regulate activity-dependent AMPA receptor recycling via exocyst complex interactions. Proc Natl Acad Sci U S A 107, 19038–19043 (2010). 10.1073/pnas.1013494107

75 Siddiqui, T. J. et al. An LRRTM4-HSPG complex mediates excitatory synapse development on dentate gyrus granule cells. Neuron 79, 680–695 (2013). 10.1016/j.neuron.2013.06.029

76 Ferreira, J. S. et al. GluN2B-Containing NMDA Receptors Regulate AMPA Receptor Traffic through Anchoring of the Synaptic Proteasome. J Neurosci 35, 8462–8479 (2015). 10.1523/JNEUROSCI.3567-14.2015

77 Farsi, Z. et al. Aberrant mRNA splicing and impaired hippocampal neurogenesis in Grin2b mutant mice.iScience 29, 114700 (2026). 10.1016/j.isci.2026.114700

78 Karataeva, A. R. et al. C-terminal interactors of the AMPA receptor auxiliary subunit Shisa9. PLoS One 9, e87360 (2014). 10.1371/journal.pone.0087360

79 Chen, X., Aslam, M., Gollisch, T., Allen, K. & von Engelhardt, J. CKAMP44 modulates integration of visual inputs in the lateral geniculate nucleus. Nat Commun 9, 261 (2018). 10.1038/s41467-017-02415-1

80 Barnes, S. A. et al. Non-ionotropic signaling through the NMDA receptor GluN2B carboxy-terminal domain drives dendritic spine plasticity and reverses fragile X phenotypes. Cell Rep 44, 115311 (2025). 10.1016/j.celrep.2025.115311

81 Wang, Y. N. et al. Controlling of glutamate release by neuregulin3 via inhibiting the assembly of the SNARE complex. Proc Natl Acad Sci U S A 115, 2508–2513 (2018). 10.1073/pnas.1716322115

82 Courtney, K. C. et al. Synaptotagmin 1 oligomerization via the juxtamembrane linker regulates spontaneous and evoked neurotransmitter release. Proc Natl Acad Sci U S A 118 (2021). 10.1073/pnas.2113859118

83 Kang, M. et al. Grin2b-mutant mice exhibit heightened remote fear via suppressed extinction and chronic amygdalar synaptic and neuronal dysfunction. Science Advances 11, eadr7691 10.1126/sciadv.adr7691

84 Sohal, V. S. & Rubenstein, J. L. R. Excitation-inhibition balance as a framework for investigating mechanisms in neuropsychiatric disorders. Mol Psychiatry 24, 1248–1257 (2019). 10.1038/s41380-019-0426-0

85 Green, S. A., Hernandez, L., Bookheimer, S. Y. & Dapretto, M. Reduced modulation of thalamocortical connectivity during exposure to sensory stimuli in ASD. Autism Res 10, 801–809 (2017). 10.1002/aur.1726

86 Wood, E. T. et al. Sensory over-responsivity is related to GABAergic inhibition in thalamocortical circuits. Transl Psychiatry 11, 39 (2021). 10.1038/s41398-020-01154-0

87 Hacohen-Kleiman, G. et al. Activity-dependent neuroprotective protein deficiency models synaptic and developmental phenotypes of autism-like syndrome. J Clin Invest 128, 4956–4969 (2018). 10.1172/JCI98199

88 Conrow-Graham, M. et al. A convergent mechanism of high risk factors ADNP and POGZ in neurodevelopmental disorders. Brain 145, 3250–3263 (2022). 10.1093/brain/awac152

89 Karmon, G. et al. Novel ADNP Syndrome Mice Reveal Dramatic Sex-Specific Peripheral Gene Expression With Brain Synaptic and Tau Pathologies. Biol Psychiatry 92, 81–95 (2022). 10.1016/j.biopsych.2021.09.018

90 Lin, C. H., Ren, Y., Tam, K. W., Conrow-Graham, M. & Yan, Z. Synaptic Deficits in Adnp-Mutant Mice Are Ameliorated by Histone Demethylase LSD1 Inhibition. Autism Res 18, 1342–1355 (2025). 10.1002/aur.70069

91 Holt, C. E., Martin, K. C. & Schuman, E. M. Local translation in neurons: visualization and function. Nat Struct Mol Biol 26, 557–566 (2019). 10.1038/s41594-019-0263-5

92 Morgens, D. W. et al. Genome-scale measurement of off-target activity using Cas9 toxicity in high-throughput screens. Nat Commun 8, 15178 (2017). 10.1038/ncomms15178

93 Joung, J. et al. Genome-scale CRISPR-Cas9 knockout and transcriptional activation screening. Nat Protoc 12, 828–863 (2017). 10.1038/nprot.2017.016

94 Zhao, Y. et al. A one-step tRNA-CRISPR system for genome-wide genetic interaction mapping in mammalian cells. Sci Rep 9, 14499 (2019). 10.1038/s41598-019-51090-3

95 Viswanathan, S. et al. High-performance probes for light and electron microscopy. Nat Methods 12, 568–576 (2015). 10.1038/nmeth.3365

96 Shaner, N. C. et al. Improved monomeric red, orange and yellow fluorescent proteins derived from Discosoma sp. red fluorescent protein. Nat Biotechnol 22, 1567–1572 (2004). 10.1038/nbt1037

97 Park, Y. G. et al. Protection of tissue physicochemical properties using polyfunctional crosslinkers. Nat Biotechnol (2018). 10.1038/nbt.4281

98 Langfelder, P. & Zhang, B. dynamicTreeCut: Methods for Detection of Clusters in Hierarchical Clustering Dendrograms. (2016). 10.32614/CRAN.package.dynamicTreeCut

99 R: A language and environment for statistical computing. R Foundation for Statistical Computing (Vienna, Austria, 2018).

100 Meyer, D., Dimitriadou, E., Hornik, K., Weingessel, A. & Leisch, F. e1071: Misc Functions of the Department of Statistics, Probability Theory Group (Formerly: E1071), TU Wien. (2025). 10.32614/CRAN.package.e1071

101 Kuznetsova, A., Brockhoff, P. B. & Christensen, R. H. B. lmerTest Package: Tests in Linear Mixed Effects Models. Journal of Statistical Software 82 (2017). 10.18637/jss.v082.i13

102 DeBruine, Z. J., Pospisilik, J. A. & Triche, T. J. (2024). 10.1101/2021.09.01.458620

103 Schneider, C. A., Rasband, W. S. & Eliceiri, K. W. NIH Image to ImageJ: 25 years of image analysis. Nat Methods 9, 671–675 (2012). 10.1038/nmeth.2089

104 Hao, Y. et al. Integrated analysis of multimodal single-cell data. Cell 184, 3573–3587 e3529 (2021). 10.1016/j.cell.2021.04.048

105 Love, M. I., Huber, W. & Anders, S. Moderated estimation of fold change and dispersion for RNA-seq data with DESeq2. Genome Biol 15, 550 (2014). 10.1186/s13059-014-0550-8

106 Yu, G., Wang, L. G., Han, Y. & He, Q. Y. clusterProfiler: an R package for comparing biological themes among gene clusters. OMICS 16, 284–287 (2012). 10.1089/omi.2011.0118

107 Dyer, S. C. et al. Ensembl 2025. Nucleic Acids Res 53, D948–D957 (2025). 10.1093/nar/gkae1071

108 Koopmans, F. et al. SynGO: An Evidence-Based, Expert-Curated Knowledge Base for the Synapse. Neuron 103, 217–234 e214 (2019). 10.1016/j.neuron.2019.05.002

109 Zheng, G. X. et al. Massively parallel digital transcriptional profiling of single cells. Nat Commun 8, 14049 (2017). 10.1038/ncomms14049

110 Nicol, P. B. & Miller, J. W. Model-based dimensionality reduction for single-cell RNA-seq using generalized bilinear models. bioRxiv (2024). 10.1101/2023.04.21.537881

111 Bais, A. S. & Kostka, D. scds: computational annotation of doublets in single-cell RNA sequencing data. Bioinformatics 36, 1150–1158 (2020). 10.1093/bioinformatics/btz698

112 Daniel, S. F. et al. *MapMyCells*: High-performance mapping of unlabeled cell-by-gene data to reference brain taxonomies. bioRxiv, 2026.2003.2006.710160 (2026). 10.64898/2026.03.06.710160

113 Pagès, H. HDF5 datasets as array-like objects in R. (2025). 10.18129/B9.bioc.HDF5Array

114 Ahlmann-Eltze, C. & Huber, W. glmGamPoi: fitting Gamma-Poisson generalized linear models on single cell count data. Bioinformatics 36, 5701–5702 (2021). 10.1093/bioinformatics/btaa1009

115 Robinson, M. D., McCarthy, D. J. & Smyth, G. K. edgeR: a Bioconductor package for differential expression analysis of digital gene expression data. Bioinformatics 26, 139–140 (2010). 10.1093/bioinformatics/btp616

116 Strimmer, K. fdrtool: a versatile R package for estimating local and tail area-based false discovery rates. Bioinformatics 24, 1461–1462 (2008). 10.1093/bioinformatics/btn209

117 Nadig, A. et al. Transcriptome-wide analysis of differential expression in perturbation atlases. Nat Genet 57, 1228–1237 (2025). 10.1038/s41588-025-02169-3

118 Korsunsky, I. et al. Fast, sensitive and accurate integration of single-cell data with Harmony. Nat Methods 16, 1289–1296 (2019). 10.1038/s41592-019-0619-0

119 Leek, J. T., Johnson, W. E., Parker, H. S., Jaffe, A. E. & Storey, J. D. The sva package for removing batch effects and other unwanted variation in high-throughput experiments. Bioinformatics 28, 882–883 (2012). 10.1093/bioinformatics/bts034

120 Peidli, S. et al. scPerturb: harmonized single-cell perturbation data. Nat Methods 21, 531–540 (2024). 10.1038/s41592-023-02144-y

