## Supplementary figures for "Concordant transcriptional and morphological remodeling revealed by *in vivo* Perturb-CLEAR"

### Extended Data Figure 1

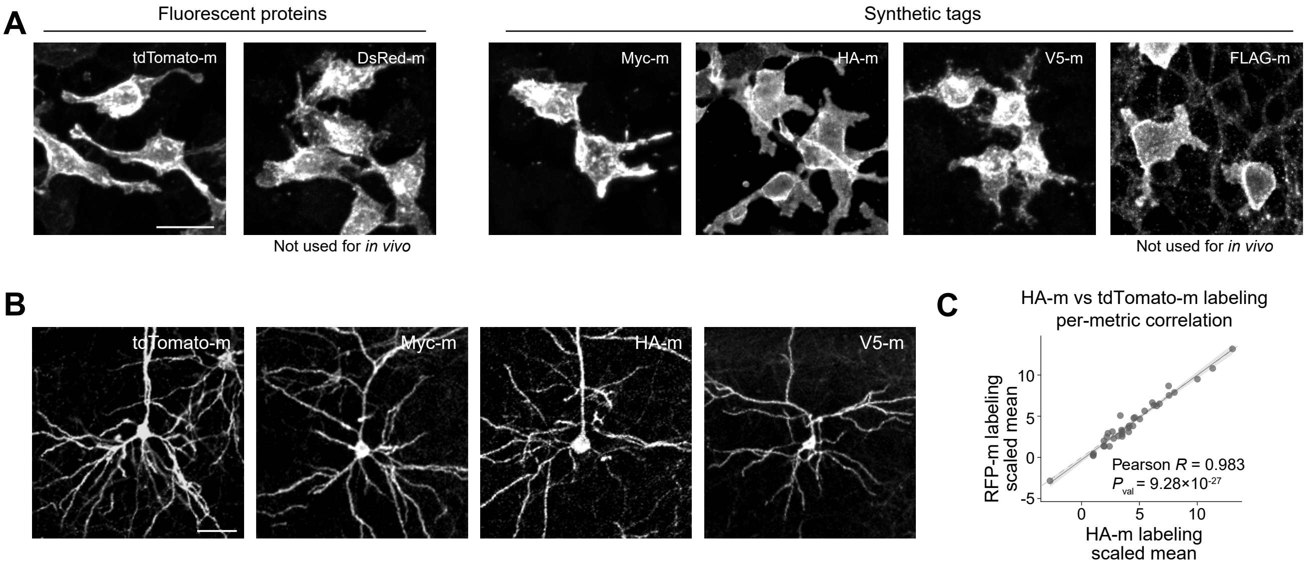

### Extended Data Figure 2

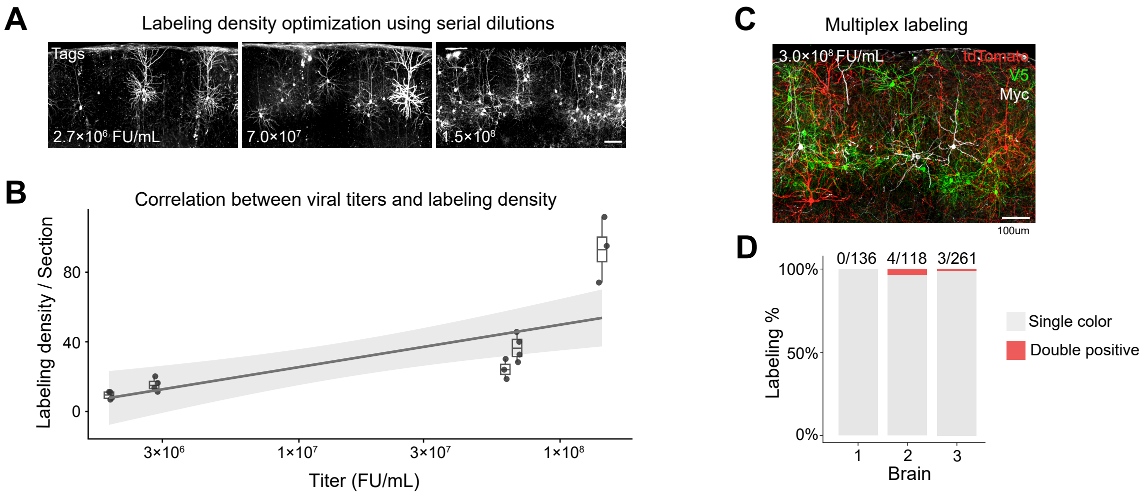

### Extended Data Figure 3

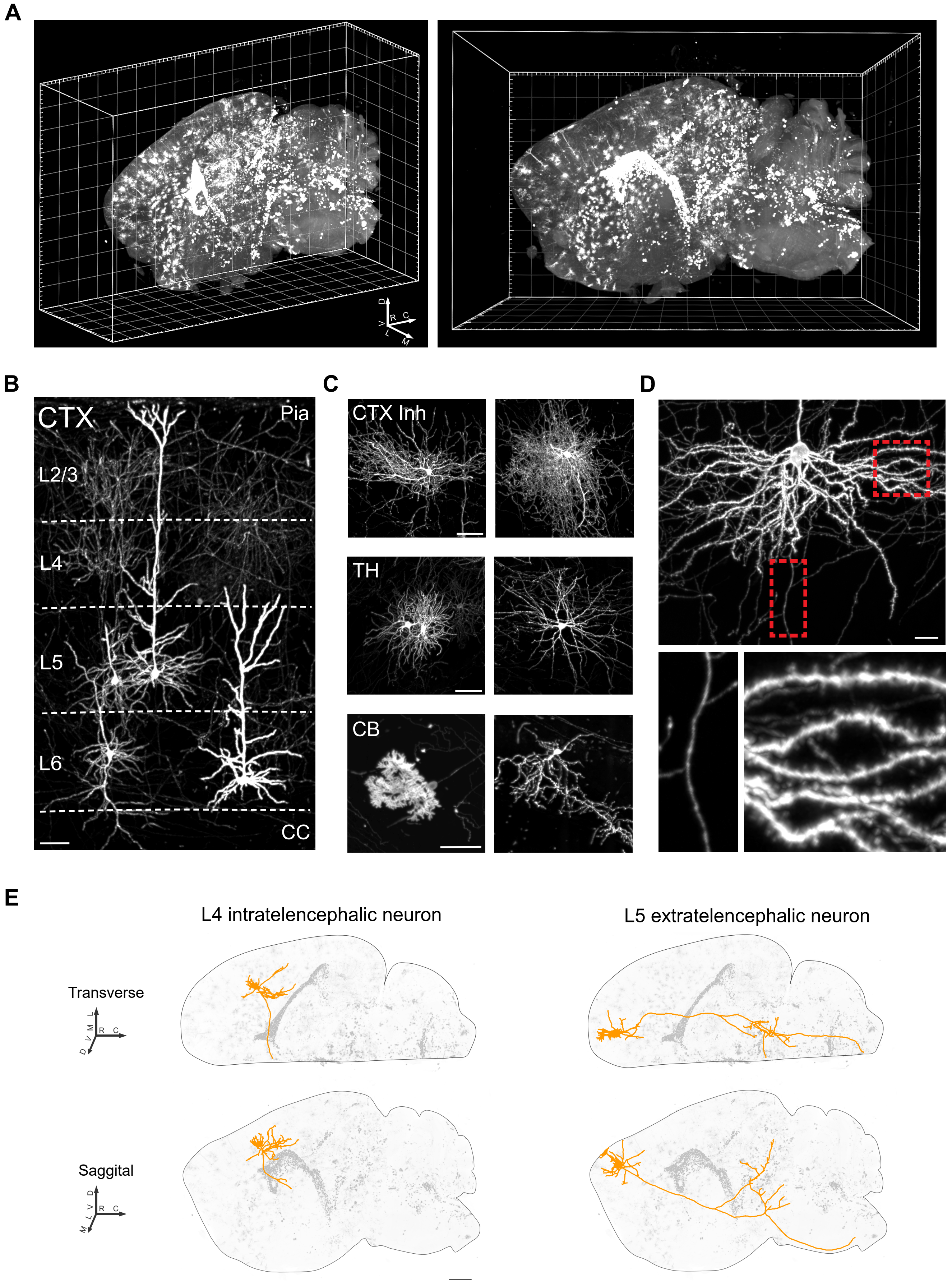

### Extended Data Figure 4

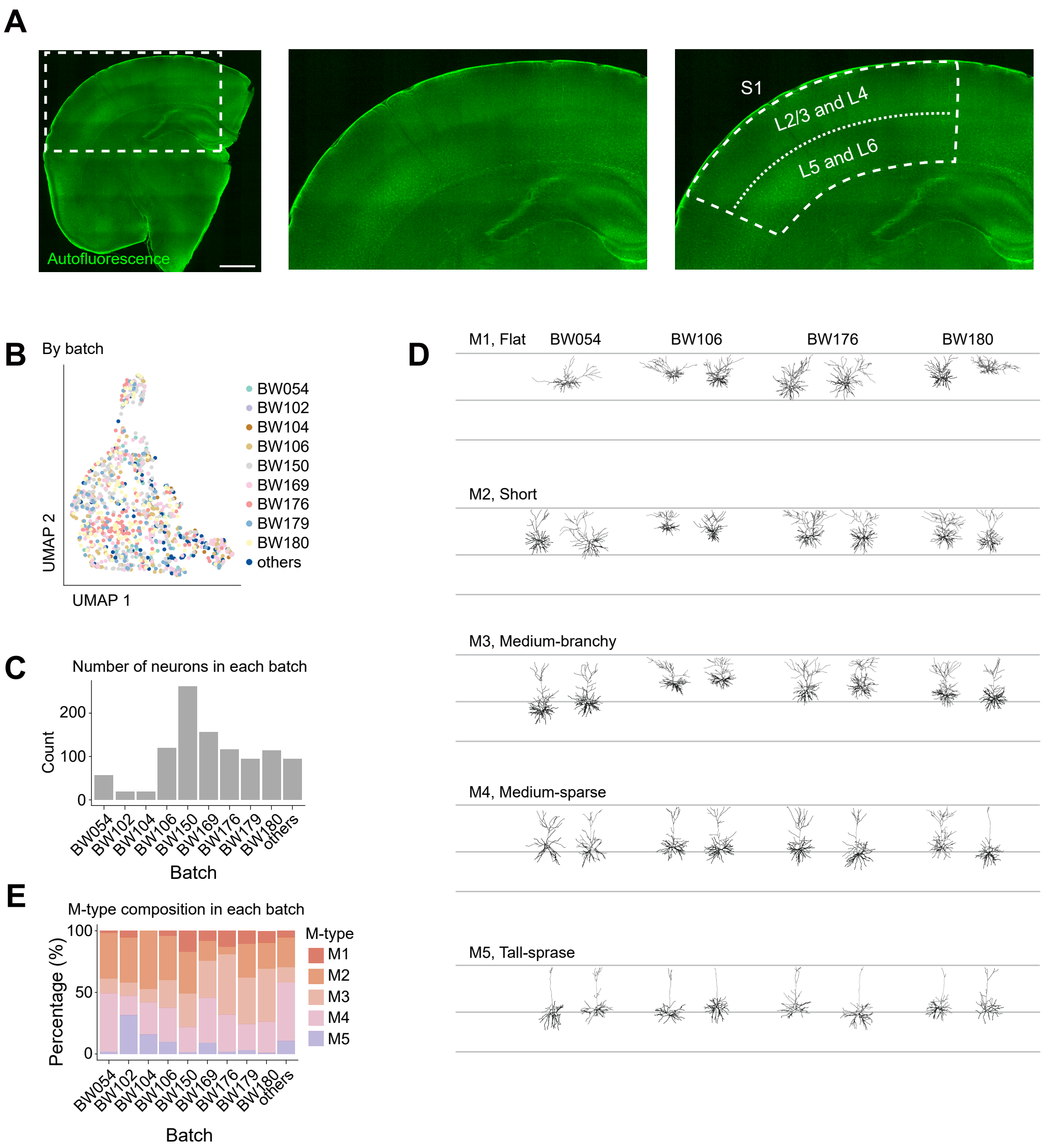

### Extended Data Figure 5

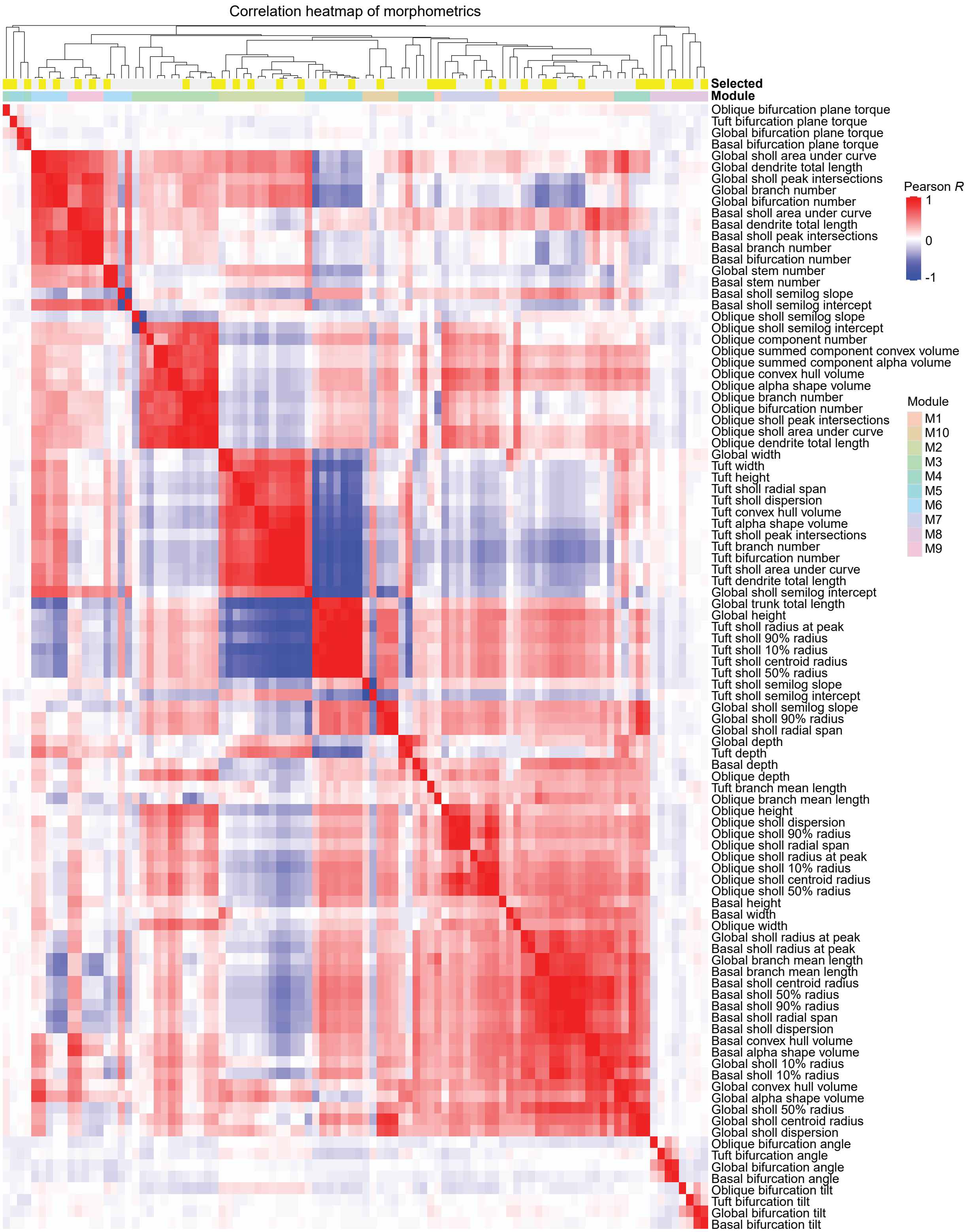

### Extended Data Figure 6

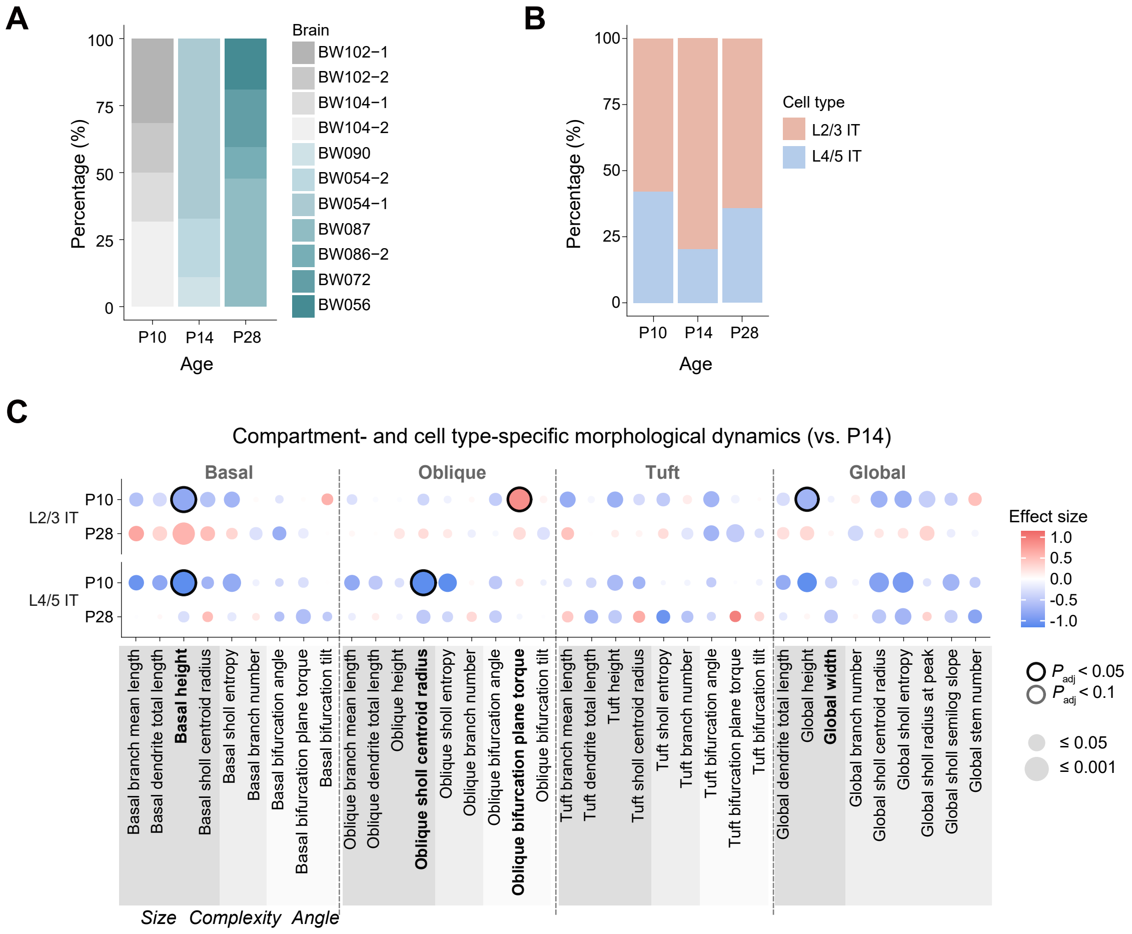

### Extended Data Figure 7

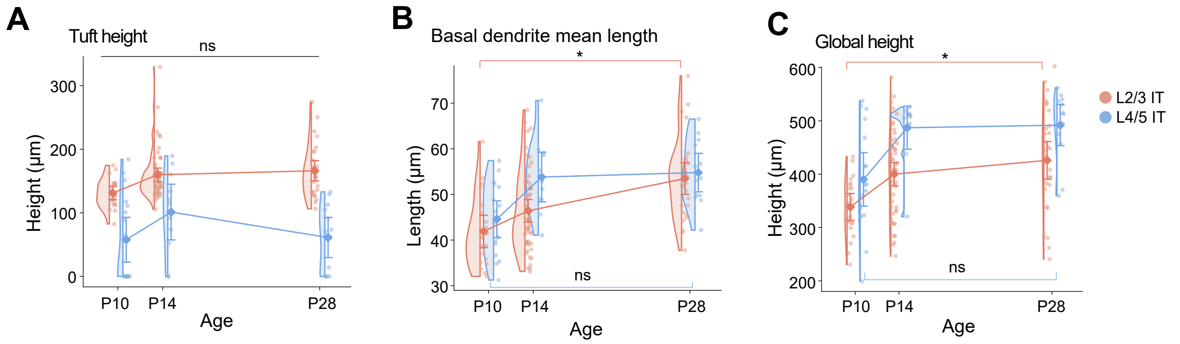

### Extended Data Figure 8

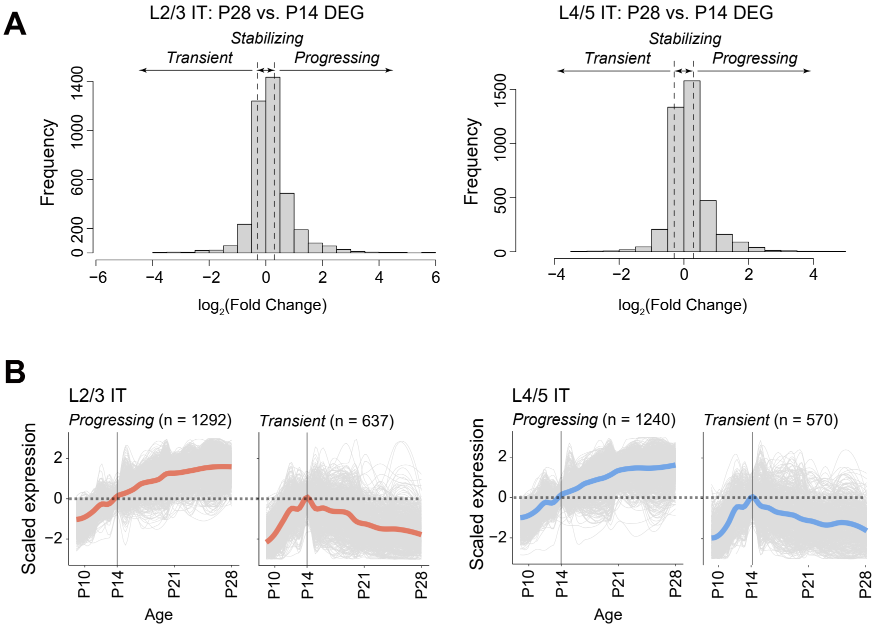

### Extended Data Figure 9

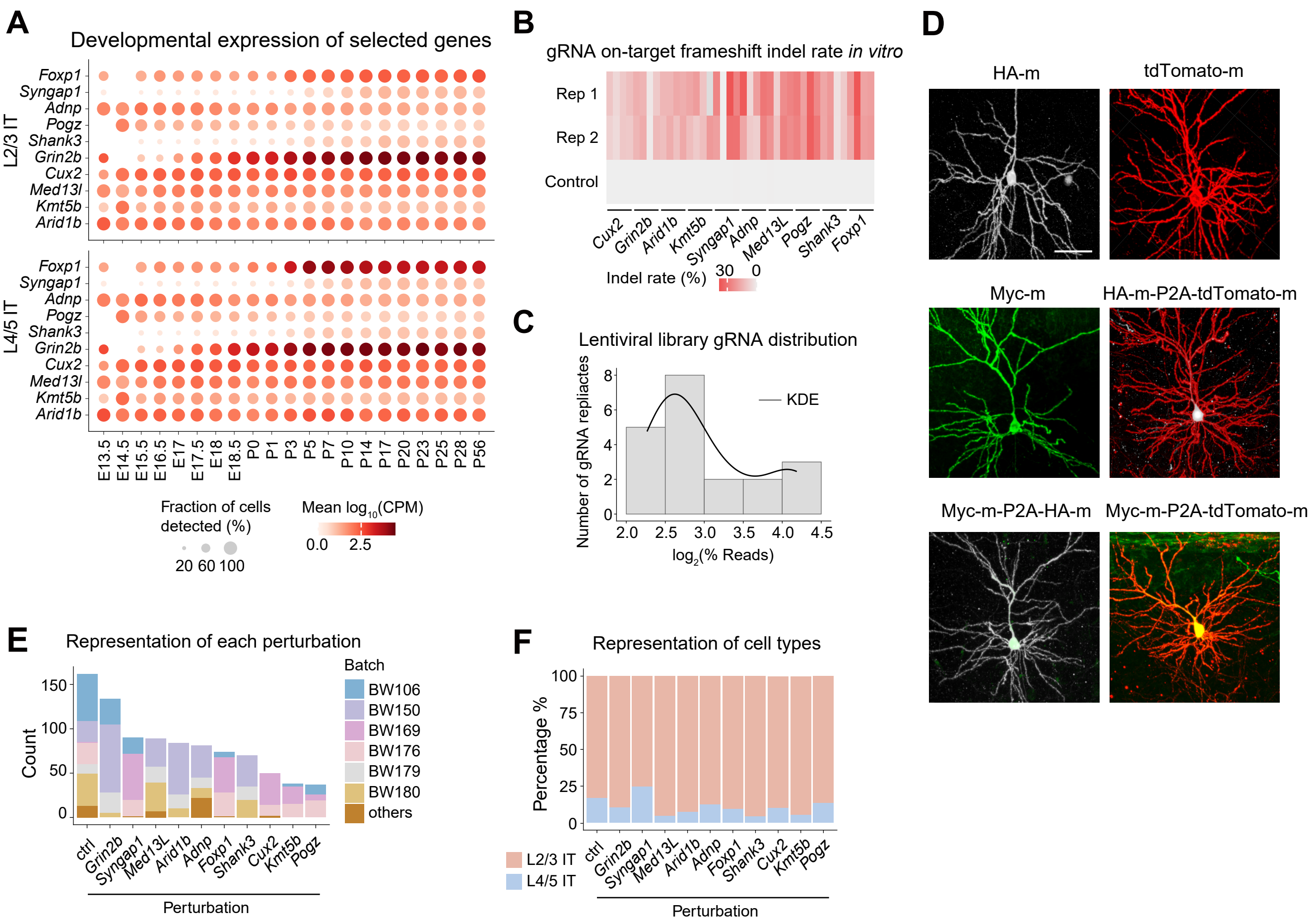

### Extended Data Figure 10

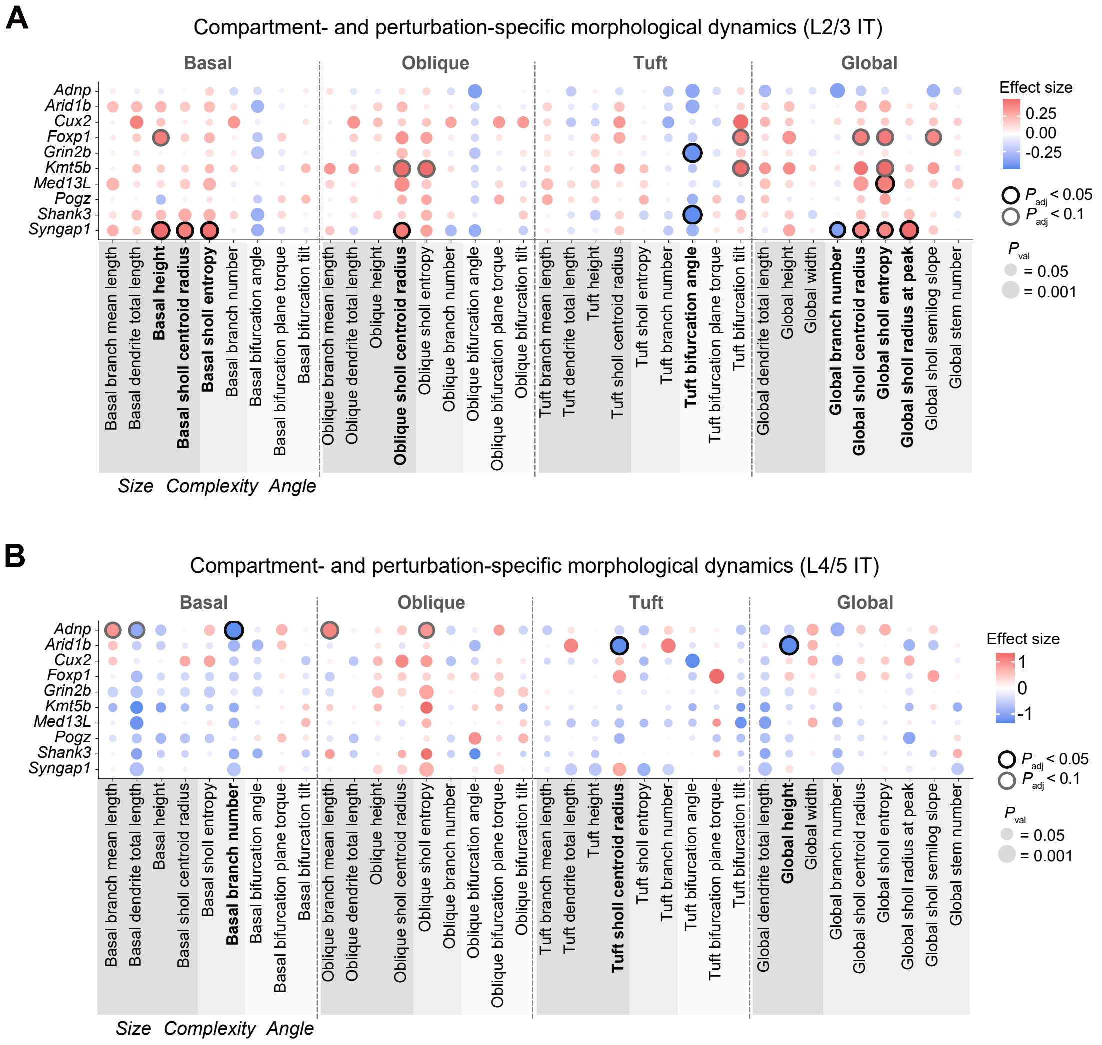

### Extended Data Figure 11

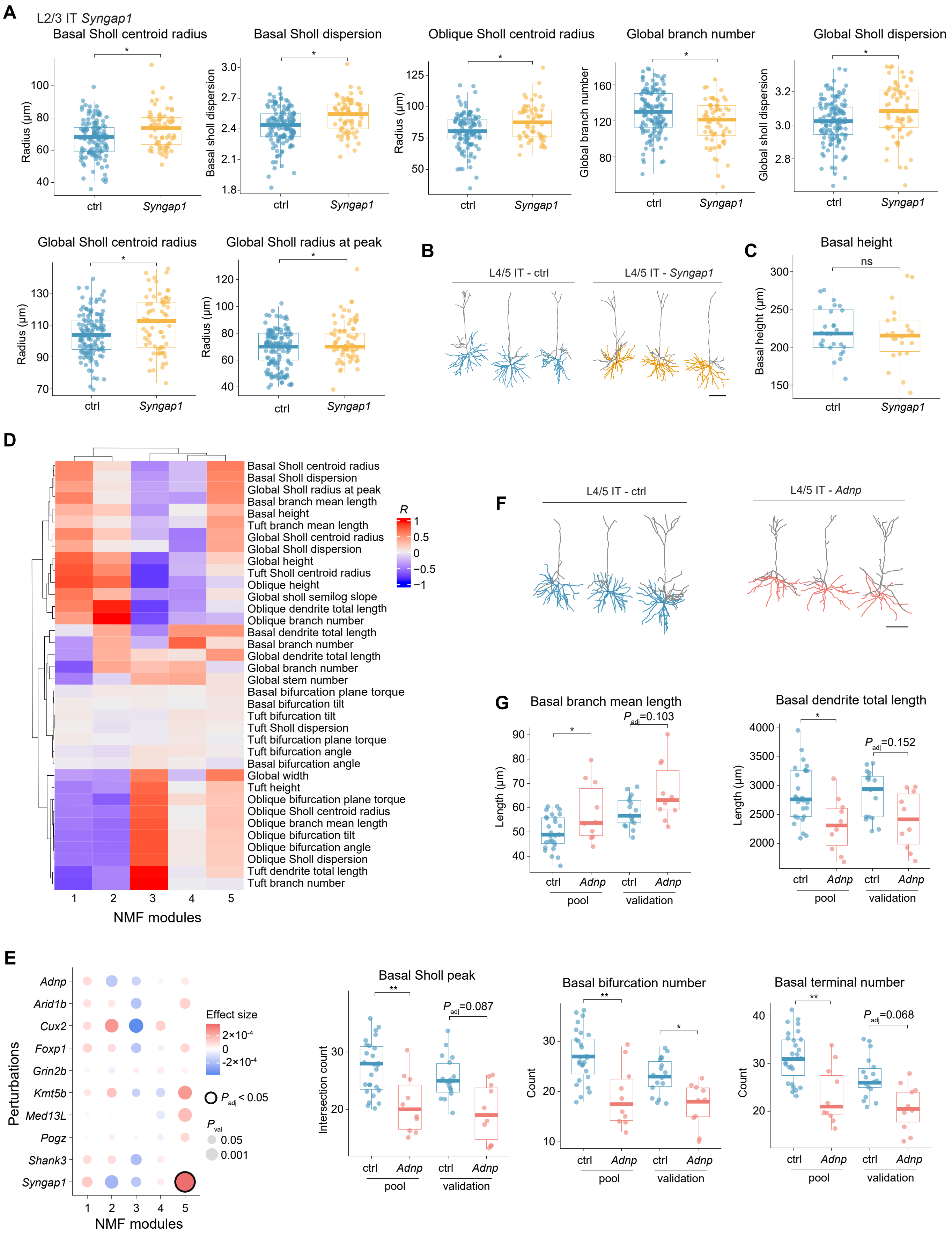

### Extended Data Figure 12

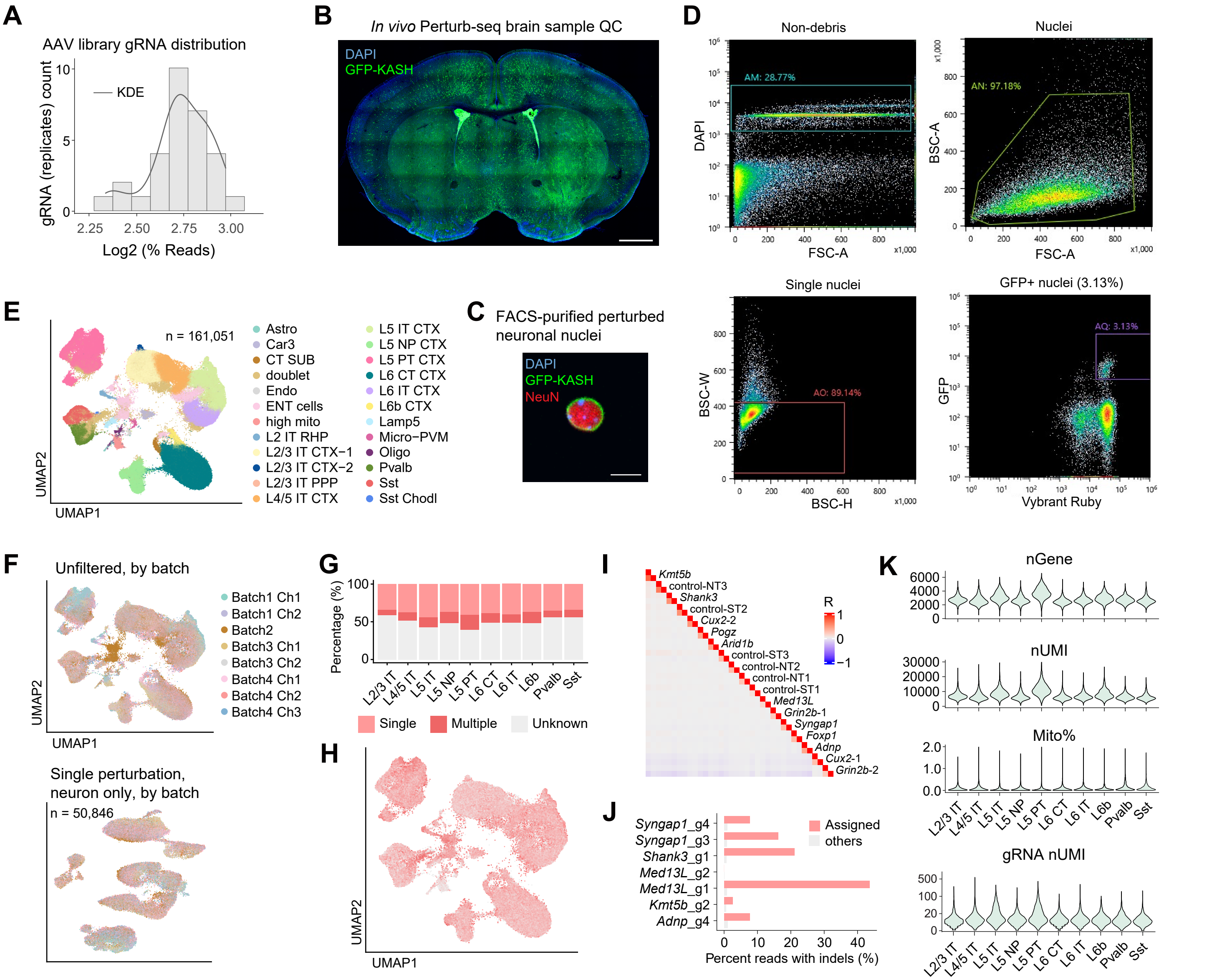

### Extended Data Figure 13

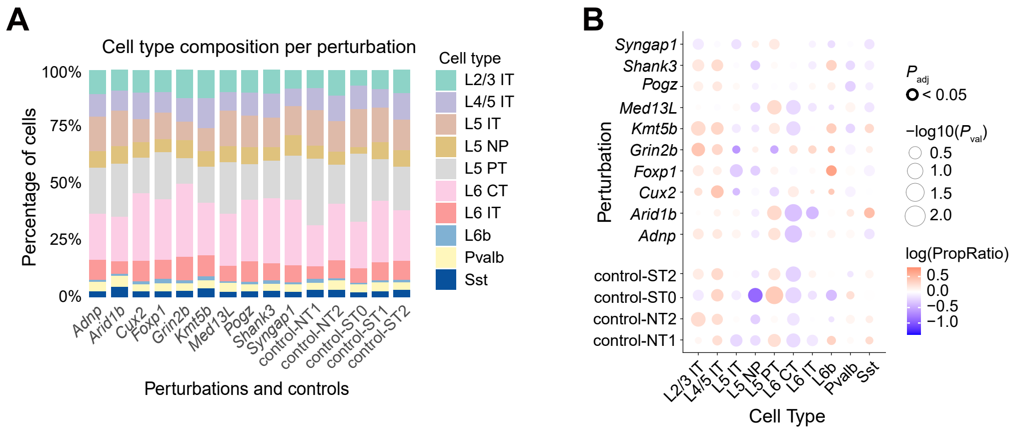

### Extended Data Figure 14

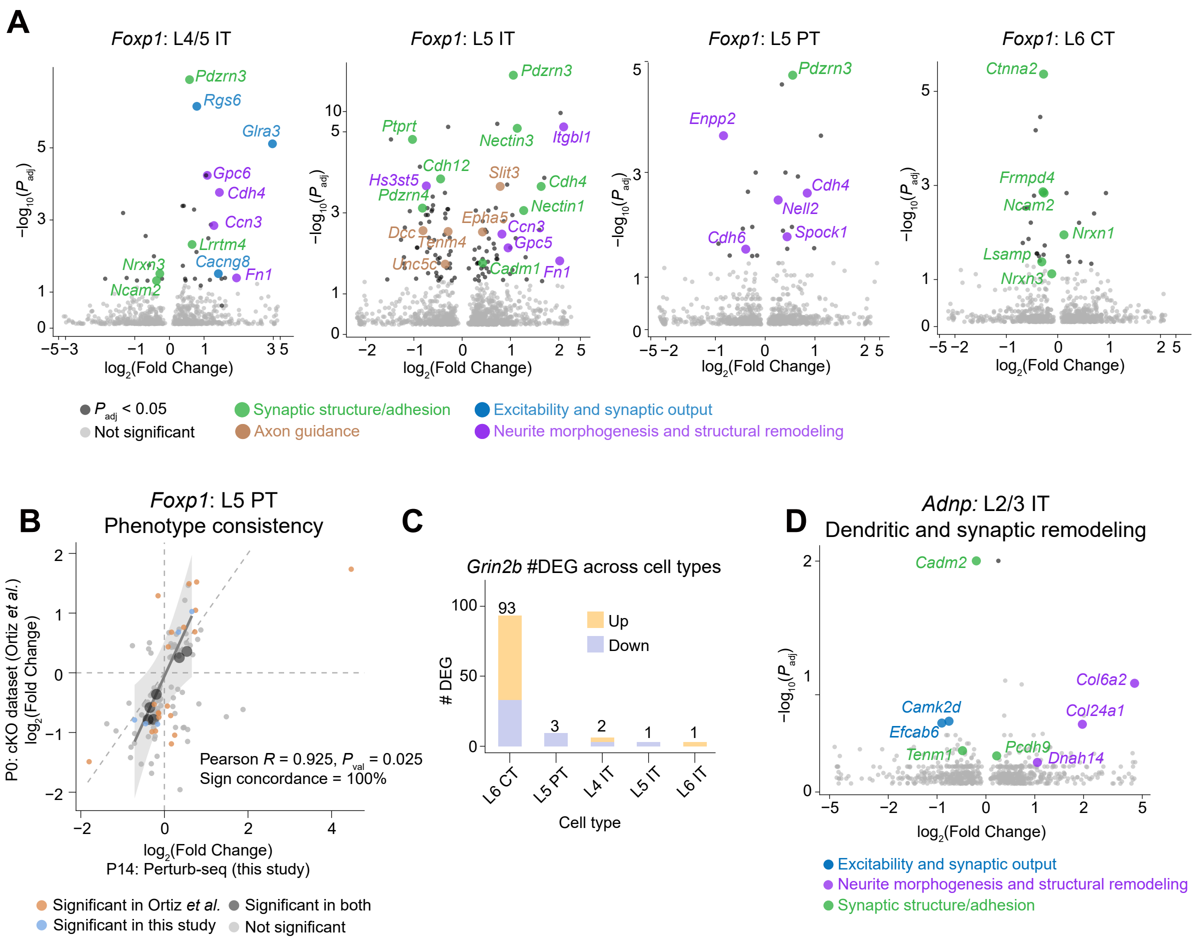

### Extended Data Figure 15

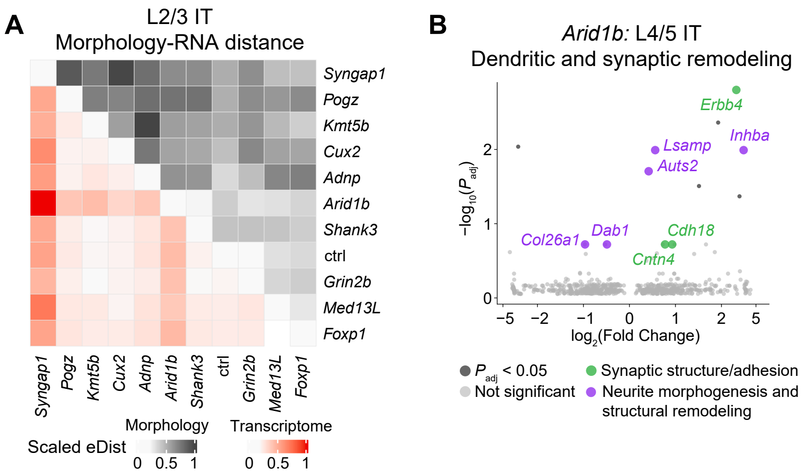
